# Direct Reconstruction of DC Cortical Conductivity from Large-Scale Electron Microscopy Data

**DOI:** 10.64898/2026.03.23.713806

**Authors:** Gregory M Noetscher, Alton Miles, Bethanny Danskin, Dexuan Tang, Mark Ingersoll, Guillermo Carlo Nuñez Ponasso, Connell Paxton, Reinhold Ludwig, Edward H Burnham, Zhi-De Deng, Hanbing Lu, Konstantin Weise, Thomas R Knősche, Bruce R Rosen, Marom Bikson, Sergey N Makaroff

**Author notes:** **Correspondence:** Gregory M Noetscher. Alton Miles, Bethanny Danskin, Dexuan Tang, Mark Ingersoll, >; Guillermo Carlo Nuñez Ponasso, Connell Paxton, Reinhold Ludwig, Edward Burnham, Zhi-De Deng, Hanbing Lu, Konstantin Weise, Thomas Knoesche, Bruce Rosen, Marom Bikson, Sergey N Makaroff. **Declaration of Interests:** The authors declare no competing interests. **Data Availability:** The numerical three-dimensional datasets described herein are available online via Tensor Conductivity Maps – ‘Mesoscale Cortical Conductivity at 50 µm Resolution’. 02/02/2026.*GitHub repository*: https://github.com/dexuantang/Mesoscale-Cortical-Conductivity-at-50-m-Resolution.

## Abstract

Electrical conductivity of cortical gray matter governs the magnitude and spatial distribution of electric fields generated by brain stimulation and intrinsic neuronal activity measured with M/EEG and intracortical recordings. However, reported macroscopic conductivity values vary by more than threefold, limiting the fidelity of bioelectromagnetic models and leaving unresolved whether this variability reflects measurement uncertainty or genuine structural heterogeneity of cortical tissue.

Here, we present a multiscale computational framework that, for the first time, attempts to derive mesoscale conductivity maps of mouse visual cortex at 50-µm resolution directly from large-volume, segmented nanometer-scale electron microscopy data. The *Minnie 65* subvolume of the MICrONS dataset is accurately subdivided into 1,224 50-µm cubic blocks. Each block contains, on average, 40-50 million membrane facets of a highly convoluted and dense cellular structure. Three orthogonal electrode pairs are applied to each isolated block to estimate the three principal components of the conductivity tensor. Quasistatic electric modeling is enabled by an iterative boundary-element fast multipole method (BEM-FMM) under the approximation of non-conducting membranes (DC conductivity).

Spatially averaged conductivity values predicted by our framework agree well with prior low-resolution measurements in rats, validating the approach. At the same time, the resulting mesoscale maps reveal pronounced conductivity granularity at 50-100 µm scales as well as significant variations in both radial and tangential directions. These results indicate that mesoscale conductivity heterogeneity could be an intrinsic structural property of the cortex. Limitations and extensions of this study are discussed in detail.

## 1. Introduction

The electrical conductivity of cortical gray matter is a fundamental physical parameter governing the strength and local spatial distribution of electric fields in the cortex, either due to various forms of brain stimulation or due to neuronal activity itself, measured by EEG, MEG, and microelectrode arrays. Personalized and targeted electrotherapy for brain disorders as well as accurate EEG/MEG and microelectrode array source reconstruction methods in neurophysiology rely on these conductivity values to accurately represent brain activity and control brain stimulation.

However, current estimates of cortical conductivity are largely theoretical or indirectly inferred [1],[2],[3],[4],[5],[6],[7],[8]. Commonly quoted values have been derived from measurements of extracellular space in small tissue samples, from fitting macroscale electrode data, or from applying a debated conversion between water-diffusion MRI and electrical conductivity. As a result, published values are rather inconsistent, sometimes mutually contradictory, and may lead to different modeling outcomes. As an example, Table 1 summarizes the conductivity values reported in several well-known studies and sources [1],[2],[3],[4],[5],[6],[7], showing that macroscopic cortical conductivity can differ by more than 300%, even before considering anisotropy. This raises the question of whether and to what extent such variability has a root in the structural variability of the cortical tissue.

**Table 1.** Some estimates for low-frequency electrical conductivity of cortical gray matter in S/m from different sources to date.

| Source | I | II | III | IV | V | VI | VII |
| --- | --- | --- | --- | --- | --- | --- | --- |
| Radial conductivity across cortical depth | 0.275 | 0.733 | 0.470 | 0.82 | 0.32-0.35 | 0.17-0.27 | ~0.52 on average |
| Tangential conductivity along cortex | isotropic | 0.231 | isotropic | Isotropic | 0.23-0.24 | isotropic | nearly isotropic |

Cortical conductivity (or, more generally, the full conductivity tensor) indeed varies across species, cortical regions and physiological states, in particular reflecting changes in extracellular space that occur during sleep, anesthesia, and other conditions [9],[10]. Nonetheless, one would expect that more reliable mesoscale conductivity estimates could be obtained for a mammalian brain directly from first principles – from the realistic, physical brain volume at nanometer resolution – rather than relying on partially indirect or phenomenological methods.

The present study is an attempt to address this gap, deriving 50 µm-resolution cortical conductivity models directly from a large-volume cellular microstructure of mouse primary visual cortex. To achieve this, two advances are prerequisite and integrated: a) serial electron microscopy petasize images of mm scale but nanometer resolution of brain parenchyma (massive datasets) and their cellular segmentation – the MICrONS dataset [11],[12],[13],[14]; b) computational method – novel boundary element fast multipole method (BEM-FMM) – to simulate current flow through such samples [15],[16],[17],[18]. While these processes require exceptional image processing and computational time, we transform these results into a conductivity map for 1,224 (17×9×8) voxels – 50×50×50 μm cubical blocks spanning layers 2/3, 4, and 5, each containing highly tortuous neuronal arbor, somas, other cells and capillaries.

The MICrONS dataset and the comparably sized human H01 dataset [19] are currently unique. However, the rapid advances in AI-assisted cellular segmentation make it reasonable to anticipate that additional comprehensive, cell-resolved brain datasets will soon become available. These future datasets are expected to span multiple brain regions and to provide exhaustive, *near-complete* labeling of neuronal and non-neuronal cellular elements.

The study is organized as follows. Section II (Materials and Methods) briefly describes the cellular content of the MICrONS 1-mm³ dataset [11],[12]. Next, it specifies a procedure of separating its largest part – *Minnie65* – into 50 μm blocks – cubical voxels containing watertight 2-manifold membrane meshes. This procedure is a central precondition for accurate numerical analysis; it is nontrivial. After that, we analyze the amount of extracellular space in every block, which includes true extracellular space plus some significant unlabeled volume. This segmentation omission reflects current limitations of large-scale automated segmentation, rather than biological absence. It needs to be corrected as described in the same section. Further, we describe the application of the boundary element fast multipole method – with three pairs of voltage electrodes attached to each block – aimed at extracting the three principal components of the conductivity tensor.

Section III (Results) provides detailed two-dimensional conductivity maps of the cortical sample overlapped with the cellular structure at mesoscale. Next, it gives complete three-dimensional conductivity maps. After that, we average the results to obtain conductivity data layerwise and when traversing across (radial conductivity) or along (tangential conductivity) the sample.

Section IV (Discussion) starts with method validation – comparison of our averaged results with biologically closest, layer-by-layer *in-vitro* multi-site LFP recordings in rodents [5], which demonstrates a nearly perfect agreement for the radial conductivity. Further, it discusses a key finding of this study – a granular character of gray matter conductivity, with neighboring regions differing by up to 50%. This observation could at least partially explain conductivity variations observed in Table 1. Next, we discuss major limitations of the study, including the necessity for an extra extracellular space correction based on independently measured data [20],[21], a restriction to only the DC or low-frequency regime (a condition of the non-conducting membrane), and the debated effect of microcapillaries, as well as possible solutions to address these weaknesses. Section V concludes the study.

## 2. Materials and Methods

### 2.1 Cellular content of the MICrONS 1-mm³ dataset

The MICrONS Phase III dataset [11] comprises a 1-mm³ volume of mouse primary visual cortex (V1), acquired with multibeam serial electron microscopy at approximately 4×4×40 nm resolution. It is separated into two parts along the anterior-posterior direction: a larger part, *Minnie 65*, and a smaller part, *Minnie 35*. The larger part, *Minnie 65*, which is shown in Fig. 1a is chosen as the target of this study. There, the gray dots are eight corners of a rectangular container tightly enclosing every cell. Biologically, this cortical region contains the full diversity of gray-matter cell classes, including excitatory and inhibitory neurons, astrocytes, oligodendrocytes and their myelin sheaths, microglia, endothelial cells, pericytes, and extracellular matrix. However, only a subset of these cellular elements was represented in the publicly released automated segmentation we used for this study (materialization = 1078 # current public as of 6/5/2024, version 1078 [12]).

**Fig. 1.**
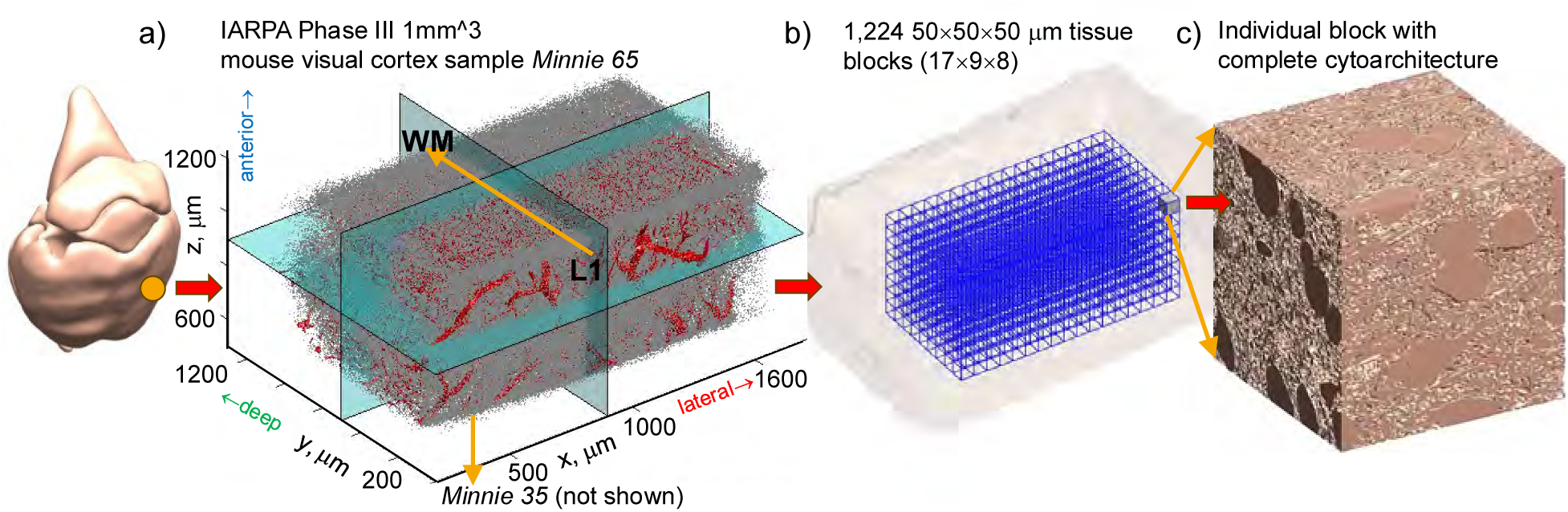
a-c) Topology of the brain volume and its subdivision. a) *Minnie 65* brain volume; the gray dots are eight corners of a rectangular container enclosing every cell; vasculature is shown in red. b) Subdivision of its central part into a set of non-overlapping subvolumes or blocks of 50×50×50 μm (the voxel size), each containing 40–50 million membrane facets. c) Individual voxel block.

Specifically, the MICrONS segmentation pipeline provided dense, high-fidelity reconstructions of neuronal cells and astrocytes [11],[13],[14] – approximately 82,000 cells in total – along with the vascular network separately segmented in [22]. Other glial populations – including oligodendrocytes, myelin sheaths, oligodendrocyte precursor cells (OPCs), and microglia -were not provided as labeled classes in the materialization used and therefore do not appear as distinct intracellular objects in downstream analyses.

This omission reflects current limitations of large-scale automated segmentation, rather than biological absence. Consequently, in this study, the intracellular compartment defined by MICrONS consists of neurons and astrocytes only, while myelin and other unlabeled glial constituents remain masked as background volume. We note that the most recent MICrONS realizations [12] are correcting this limitation.

### 2.2 Separation of convoluted brain volume into 50 μm blocks – cubical voxels

The *Minnie 65* brain volume is composed of ∼500 billion membrane facets when a resolution of 200 nm is maintained. This originally segmented (triangular membrane segmentation) brain volume and the vascular volume independently segmented in [22] were initially subjected to alpha wrapping of CGAL [23] to create smoother membrane surfaces (suitable for accurate modeling) at 200 nm resolution. The entire brain volume was further subdivided into a set of non-overlapping subvolumes of 50×50×50 μm (the voxel size) as in Fig. 1b,c, each containing 40–50 million facets. This set is so chosen to cover the ‘most reliable’ volumetric part of the sample, which is well separated from its boundaries. Fig. 1b shows the *Minnie 65* sample outline (transparent pink) with the assembly of 1,224 50-μm voxel blocks located in the ‘best’, most reliable part of the sample, which is well separated from its boundaries. The voxel blocks span L2/L3, L4, and L5. We note that every block has a margin of 1 μm on each side so that its initial size is in fact 52×52×52 μm. This is important for the mesh cutting method described below.

### 2.3 Construction of 2-manifold membrane meshes respecting block boundaries

The block subdivision procedure is nontrivial. At Step 1, we search through all triangular membrane faces of the original sample and retain only faces whose centers are within the given block volume. Every block has an overlap of 1 μm on each side to ensure that the faces intersected by the block boundaries are still included into consideration. This initial operation gives us a *raw* block mesh with holes that have irregular staircase boundaries in 3D.

At Step 2, we apply a custom mesh-plane intersection algorithm six times, one for each of the six block sides, and subdivide all triangular faces crossing the block boundaries to have the new mesh edges exactly at the boundaries. We retain the subdivided triangles within the 50×50×50-μm block volume and exclude all other subdivided triangles. The mesh so constructed (the ‘inner’ block mesh) precisely respects the block boundaries and has no faces or face pieces outside the block. The inner block mesh still has open holes at each of its sides due to cutting out the remainder of the cytoarchitecture. However, the holes now have well-defined boundaries in the form of closed 2D planar contours in their respective planes. Therefore, they can be closed.

At Step 3, a 2D Delaunay triangulation is applied to every such planar contour to ‘close the hole’ – to fill it out with facets of approximately the same size of 100 nm. When a hole is spanning two (or three for a block edge) block sides and becomes non-planar, a modified closure algorithm is applied. Closing all the holes results in an ‘outer’ mesh of the block. Finally, the mesh with the hole caps (the outer mesh) is combined with the inner mesh and all duplicated nodes are removed. As a result, we obtain a combined block mesh with consistent outer normal vectors. This block mesh is 2-manifold and is suitable for numerical modeling.

Fig. 2a,b demonstrates typical assembly of two 50×50×50 μm block meshes. All holes – which appear due to cutting out the membrane meshes – are accurately closed. After this operation, all final cellular meshes become 2-manifold. Every block is further embedded in a meshed container (shown in transparent blue) such that a gap of 1 μm always exists between the cellular structure and electrodes and/or insulating walls of the container. This is done to avoid overlapping of cell boundaries with electrodes and associated numerical instability. The container sides are also meshed.

**Fig. 2.**
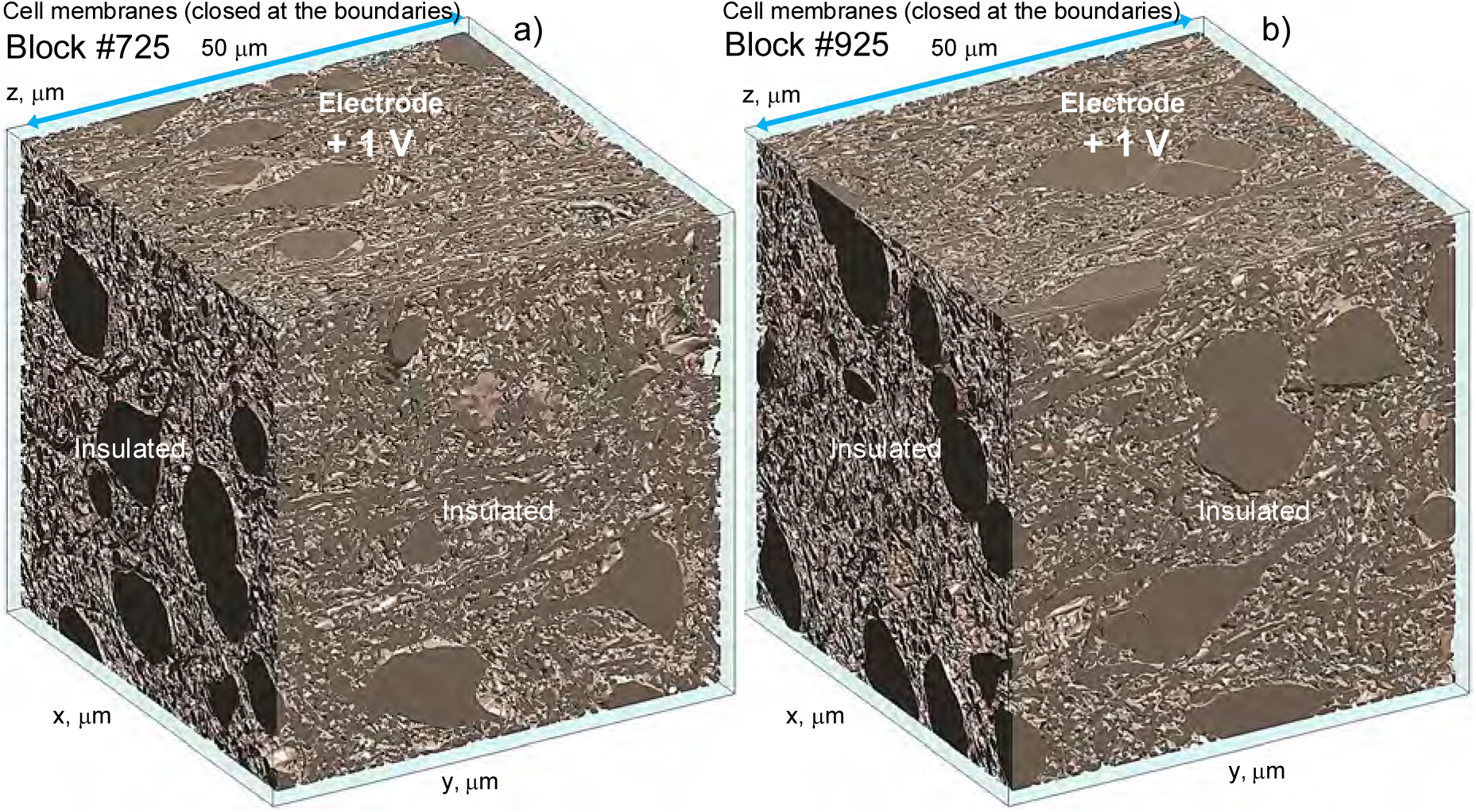
a,b) Example 50×50×50 μm block structures with all holes – which appear due to cutting out the membrane meshes – accurately closed. Every block is further included in a container shown here in transparent blue such that a gap of 1 μm always exists between the cellular structure & electrodes/insulating walls of the container.

### 2.4 Estimation of relative volume *v* occupied by neurons and astrocytes for every block

A grid of uniformly distributed 3D query points on the size *N* × *N* × *N* was introduced within every block, and an inside/outside test was performed for the cellular volume. Each block was processed independently, and the resulting inside/outside classifications were concatenated to form the final result. Blockwise evaluation was parallelized, with independent point blocks distributed across workers. It was found that the values of *N* ≥ 60 (a ∼0.8 μm query point spacing) provided results within 1% of the converge limit. A convergence test was further performed to assure that the computation error is less than 1%. Once the relative cellular volume *v* is found, the actual conducting fraction, which includes extracellular space (ECS) plus the unlabeled volume, is obtained in the form

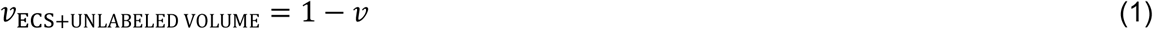

Fig. 3a depicts the effective or actual conducting fraction, *v*_ECS+UNLABELED VOLUME_, obtained in this way. It includes extracellular fluid (ECS) plus unlabeled volume, for each of the 1,224 square blocks shown by different colors of labeling spheres located at the center of each block. Fig. 3b gives *v*_ECS+UNLABELED VOLUME_ values averaged over the principal cross-section of the sample as a function of distance across the cortex. Dashed lines show two margins corresponding to maximum/minimum values for the given sample cross-section. Table 2 below summarizes average and extreme values of the conducting fraction for different layers. Averaging is made for every radial sample cross-section (9 such cross-sections in total) spanning 17×8 blocks in the medial-lateral and anterior-posterior directions. The values from Table 2 are in line with the original MICrONS estimates [13],[14],[11], which predict 0.5-0.65 neuronal fraction and 0.05-0.1 astrocytes fraction – this gives us 0.25-0.45 of the remaining *v*_ECS+UNLABELED VOLUME_. On the other hand, the actual biological fraction of *v*_ECS_ is estimated as 0.15-0.25 [20],[21],[24],[25],[26]. This means that the corresponding correction should be made.

**Fig. 3.**
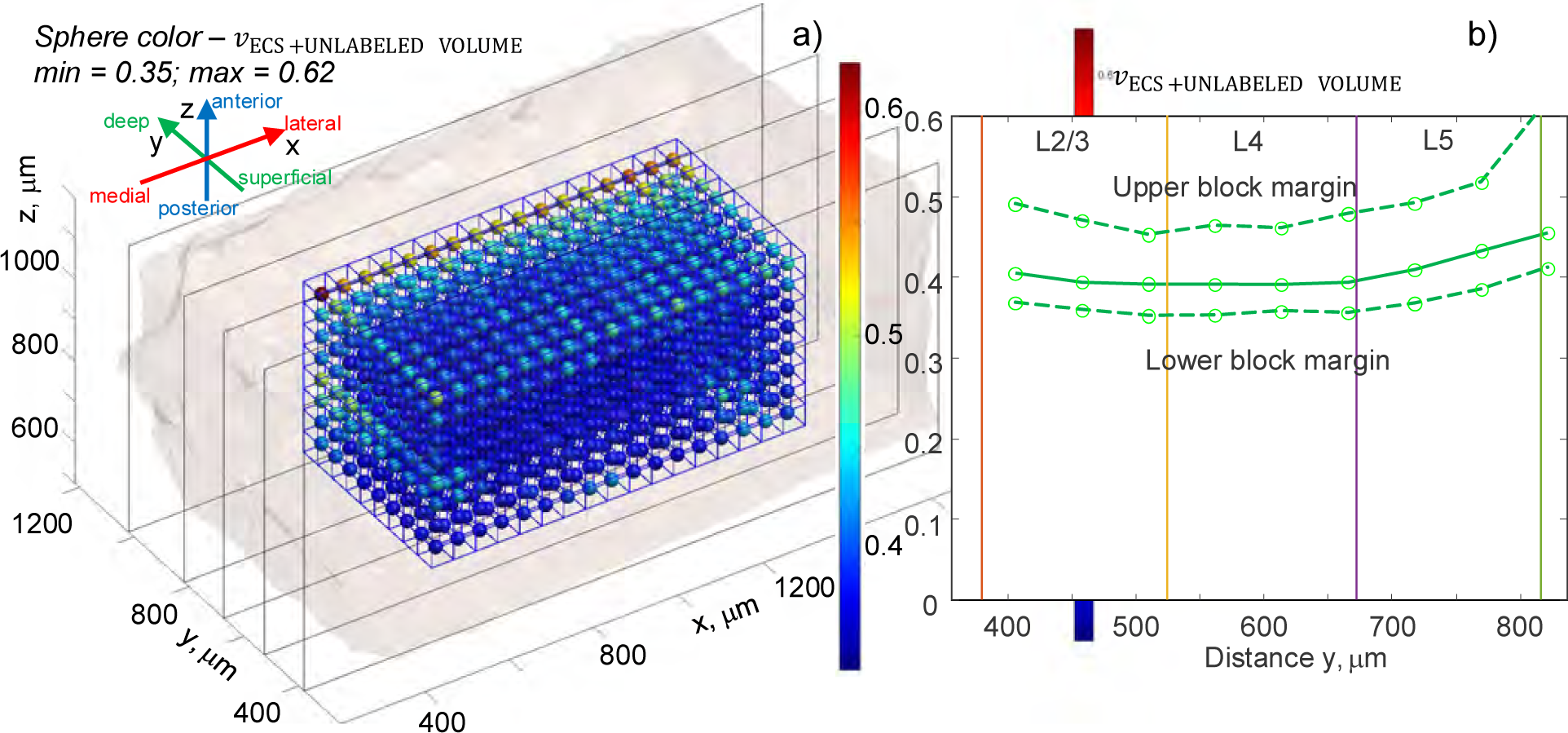
a) Effective(actual) conducting fraction *vv*_ECS+UNLABELED VOLUME_ which includes ECS plus unlabeled volume, for every of 1,224 square blocks spanning L2/L3, L4, and L5 of the *Minnie 65* MICrONS sample shown by different colors of labeling spheres located at the center of each block. b) The same values averaged over the principal cross-section of the sample as a function of distance across the cortex. Dashed lines are two margins corresponding to upper/lower values.

**Table 2.** Average and maximum/minimum values of the conducting fraction, *v*_ECS+UNLABELED VOLUME_, for different layers.

| Layer | L2/L3 | L4 | L5 | L6 |
| --- | --- | --- | --- | --- |
| $v_{\text{ECS+UNLABELED VOLUME}}$ | **0.39** | **0.39** | **0.42** | **NA** |
| Minimum block value | 0.35 | 0.35 | 0.37 | NA |
| Maximum block value | 0.49 | 0.48 | 0.62 | NA |

### 2.5 Correction of block conductivity tensor due to segmentation incompleteness

The primary MICrONS segmentation includes neuronal and astrocytic compartments but does not label myelin, oligodendrocytes, or glia. Consequently, the homogenized conductivity tensor ***σ*** computed from the segmented geometry will overestimate the true gray-matter conductivity, because part of the volume initially treated as electrically conductive extracellular space (ECS) is in fact insulating myelin/oligodendrocyte/glia cells. To account for this constrained-conductor regime, we applied a correction derived from classical effective-medium theory [27],[28],[29],[30], in which the effective macroscopic conductivity of a composite medium is proportional, to first order, to the connected conducting volume fraction when the system lies well above the percolation threshold. In such media, small variations in insulating content produce first-order changes in effective conductivity given by

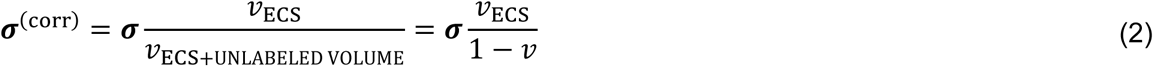

where *v*_ECS_is the true relative amount of the conducting extracellular space. This expression preserves the anisotropy of ***σ*** but rescales its magnitude by the fraction of conducting volume removed by the addition of previously unsegmented insulating material. Eq. (2) reflects the fact that, when the conductive volume fraction is limited, even a small addition of insulating material can reduce the effective conductivity disproportionately – analogous to partially occluding an already narrow conducting pathway.

For rodent’s gray matter (adult rats), measured relative values of *v*_ECS_ are available from the literature [20],[21]; the corresponding values used in this study at the layer centers follow [20],[21]:

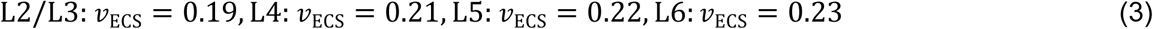

Other relevant values are obtained by interpolation. With this information, Eq. (2) becomes a complete result, with two other parameters – ***σ*** and *v* – found from the present analysis.

### 2.6 Using condition of a non-conducting membrane

The present study is using the condition of a non-conducting membrane. This is because the ohmic surface leakage conductance *G_mDC_* of idle neuronal cells is typically very small (∼1e-12 S/μm^2^) [31],[32],[33],[34] and the ohmic conductance of astrocytic membranes is not significantly larger. However, the capacitive AC conductance of the membrane *G*_*mAC*_ = *j*ω*C*_*m*_ (where *C*_*m*_∼1e-14 F/μm^2^ [33] is membrane capacitance; *ω* is angular frequency) may become quite significant when frequency exceeds approximately 1-5 kHz.

Our analysis of individual MICrONS neurons [11],[12] with permeable passive membranes shows the condition of the non-conducting membrane works well when frequency does not exceeds 100 Hz. Therefore, we restrict the present analysis to a DC conductivity tensor or to a low-frequency AC regime when the frequency does not exceed approximately 100 Hz. Also note, that in the context of brain stimulation, the condition of the non-conducting membrane also corresponds to the so called state of the initial polarization [35],[36],[37].

### 2.7 Numerical solution with boundary element fast multipole method (BEM-FMM)

For the non-conducting membrane, the intracellular and extracellular problems become decoupled, and the extracellular fields is found by solving only an exterior Neumann problem with the single-layer (layer of monopoles) charge density *c*_*e*_. On non-conducting membrane surfaces *S* with the outer normal vector ***n***, Fredholm equation of the second kind has the form [15]

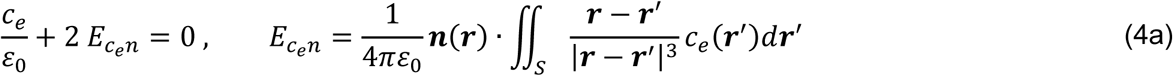

Here, *ɛ*_0_ is the dielectric constant of vacuum. Two opposite sides (say, ±*z*) of the 52×52×52 μm container with the 50×50×50 μm cubical tissue block inside are assigned voltages *V* of ±1 V where all other sides are kept insulated – the normal component of the electric field just inside the container is set to zero. On the electrode surfaces, the Fredholm equation of the first kind applies

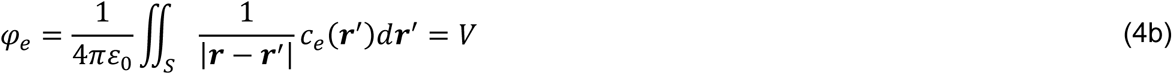

The container shown in Fig. 2a,b is further filled with extracellular fluid with an auxiliary conductivity *σ*_*e*_ = 1 S/m. The exact conductivity value matters not for the subsequent analysis. In this study, the normalized results for conductivity will be presented along with the absolute values – the former needs to be multiplied by the realistic value of the extracellular conductivity when the absolute values are concerned.

The mixed boundary value problem (4) within the enclosing cube is solved via BEM-FMM [15],[16],[17],[18]. The fast multipole method (FMM) [38],[39] allows us to solve for up to 100 million membrane facets in a block. We use a block-diagonal electrode preconditioner and employ up to 350 GMRES (Generalized Minimum Residual Method) iterations, which guarantees us the relative residual to be less than or equal to 1e-4 for ∼95% of the cubes (voxels). For the remaining voxels, a relative residual of 1e-3 is guaranteed.

The BEM-FMM solution yields the induced single-layer charge density on both membrane surfaces, which is illustrated in Fig. 4a,b, as well as the container boundaries, including the electrodes themselves. Further, the electric field everywhere in the extracellular volume within the cube was computed via Coulomb’s law, again using the fast multipole method. The typical electric field distribution in a 2D cross-section of two representative blocks is illustrated in Fig. 5b. Here, magnitude of the total electric field is plotted in a 2D cross-section of the container; its position is shown in the inset of Fig. 5a. In Fig. 5b, color encoding corresponding to the equivalent homogeneous solution is marked by an arrow on the colorbar. This problem is linear; it can be scaled proportionally to any value of the applied electrode voltage.

**Fig. 4.**
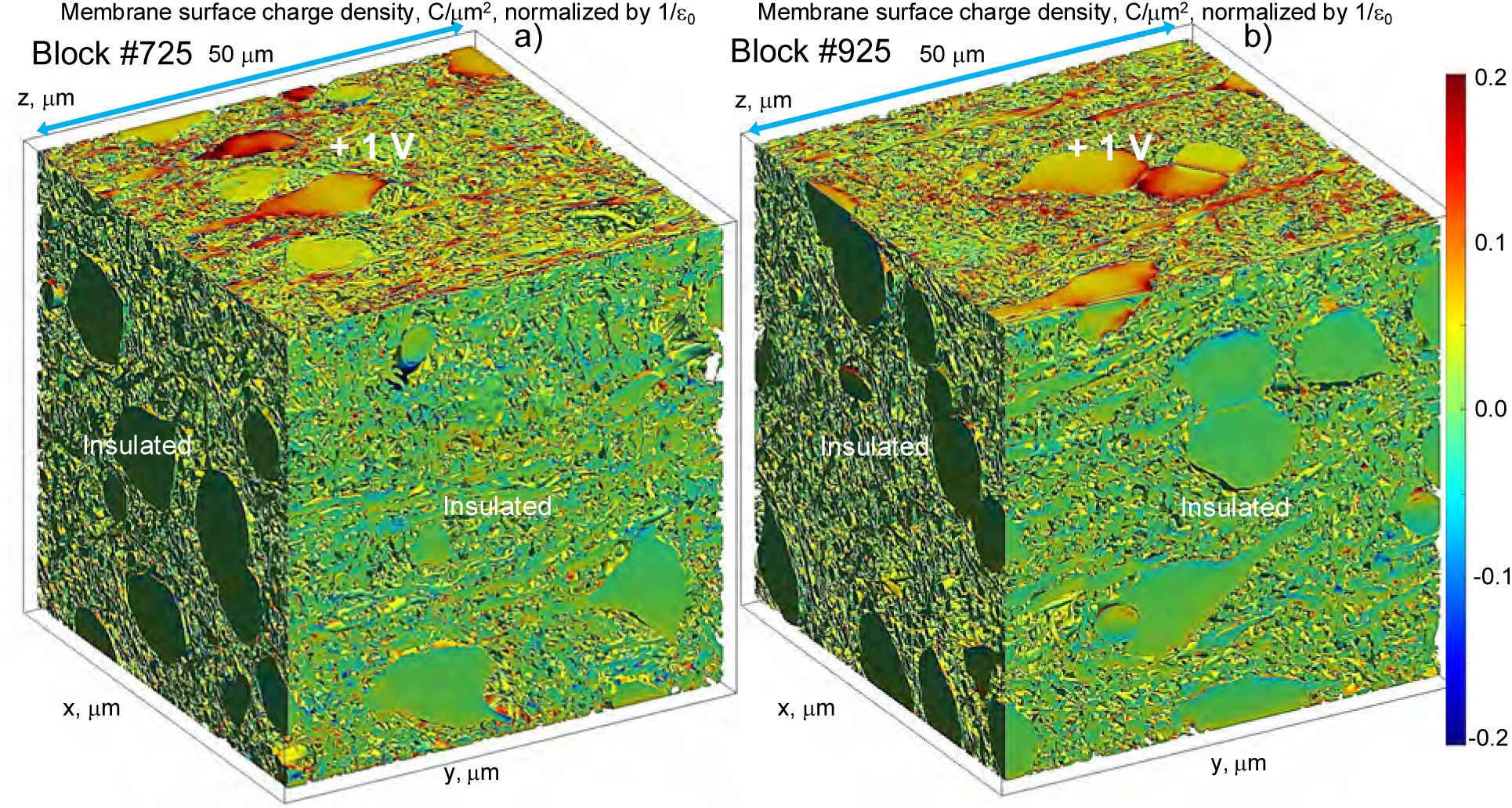
Induced membrane charge distribution for two blocks from Fig. 2 when a ±1 V voltage is applied to a pair on the opposite walls of the container while all others are kept insulated obtained with BEM-FMM. Finding electrode electric fields gives us the total electrode current.

**Fig. 5.**
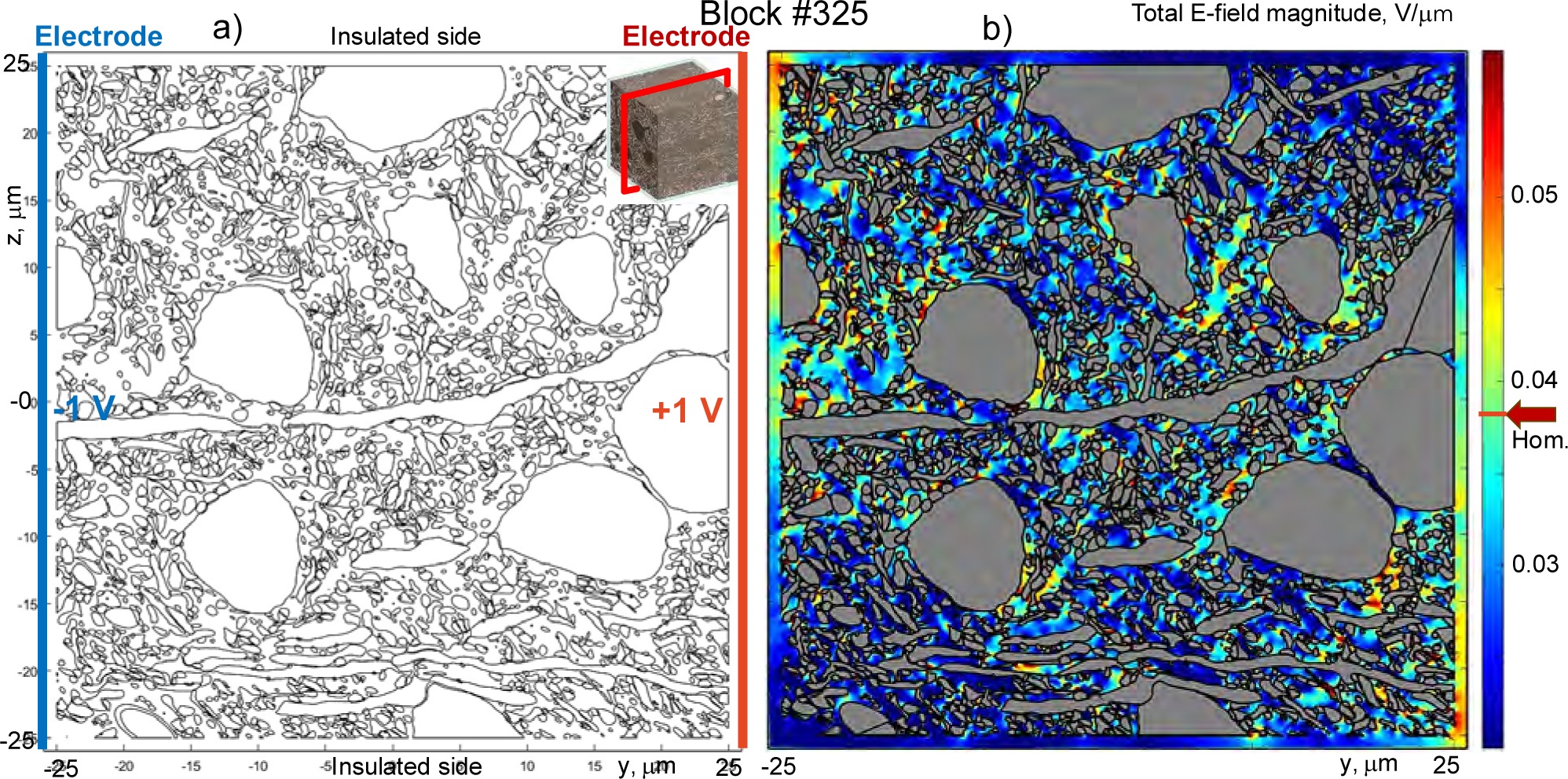
Typical electric field distribution in a 2D cross-section of a representative block when a voltage of ±1 V is applied to the container sides. The cross-section passes through block center. a) Membrane topography and cross-section position (inset on top right). b) Magnitude of the total electric field is plotted in a 2D cross-section of the container. Color encoding corresponding to the equivalent homogeneous solution is marked by an arrow on the colorbar.

The local current density at the electrode surface was finally obtained as the electric field just inside the electrode surfaces multiplied by the extracellular conductivity *σ*_*e*_ since the medium in direct contact with electrodes is always homogeneous – the container is larger than the enclosed tissue block by 1 μm. The net electrode current was obtained as an integral of the current density over the electrode surface. The current balance (equality of net inflowing and outflowing currents) was controlled during computations; the corresponding deviation did not exceed 0.1% for ∼95% of all cases and 1% otherwise.

### 2.8 Construction of block conductivity tensor

To obtain the net current in all three principal directions (*x*, *y*, *z*), the numerical solution for each container was repeated three times, for three different pairs of voltage electrodes. The following method was used to extract the block conductivity tensor. The total resistance of the container in one direction is the voltage difference of 2 V divided by the net electrode current. This total resistance is approximately but not exactly equal to the true resistance of the 50×50×50 μm block, *R*_*V*_.To find *R*_*V*_ more accurately, we represent the total container resistance as a series combination of (i) a parallel combination of the true resistance of the block and equal resistances of four narrow 1 μm wide gaps filled with extracellular fluid and adjacent to the four insulating container sides and; (ii) two equal resistances of the 1 μm wide gaps adjacent to the electrodes themselves. Knowing the rectangular gap geometry, we then find the true *R*_*V*_. Finally, the ratio 1/(50*R*_*V*_) gives us the absolute block conductivity in S/μm which is normalized to the conductivity of the homogeneous extracellular space (an auxiliary value *σ*_*e*_ = 1 S/m = 10^−6^ S/μm is used in this study) and is denoted by *σ*_*i*_ where *i* = 1,2,3 (the x-, y-, and z-axes). The values *σ*_*i*_ are the desired diagonal components of the electric conductivity tensor ***σ*** for the given block.

## 3. Results

### 3.1 Two-dimensional conductivity maps

A detailed insight into the mesoscale conductivity behavior and its link to the cortical architecture is provided by two-dimensional conductivity maps given in Fig. 6 Panels I-VIII. Here, we present a) position of the observation plane within the sample when moving from anterior to posterior direction; b) all cross-sections of the original cellular brain volume containing the selected block grid in the coronal (frontal) plane obtained with an in-house method that potentially expands the capability of the commonly used Neuroglancer ; c) the corresponding conductivity maps for three different components of the conductivity tensor in the same plane which is always passing through the block centers. The two-dimensional conductivity maps in Fig. 6 Panels I-VIII gradually evolved in the direction from anterior to posterior.

**Fig. 6.**
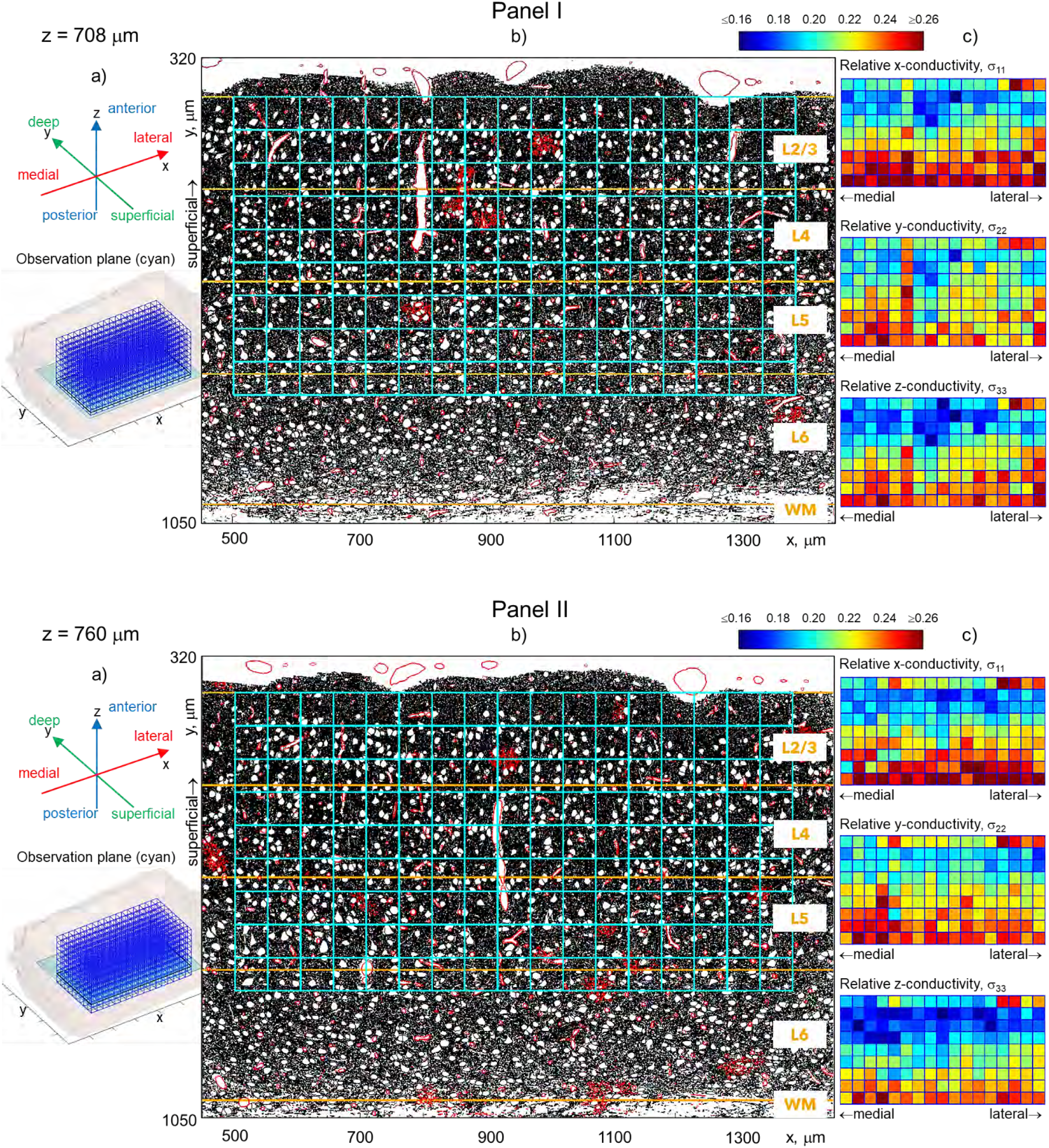
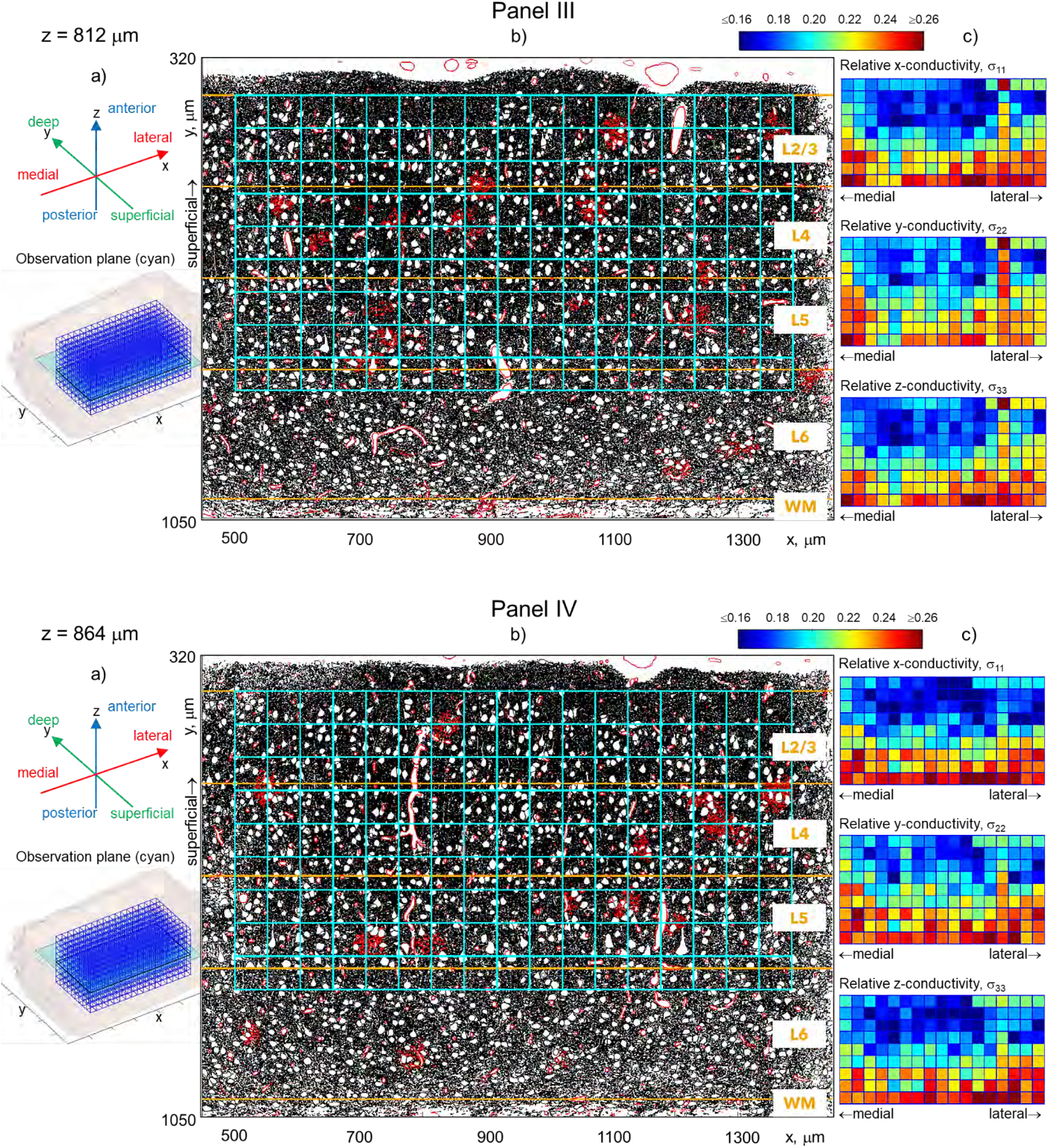
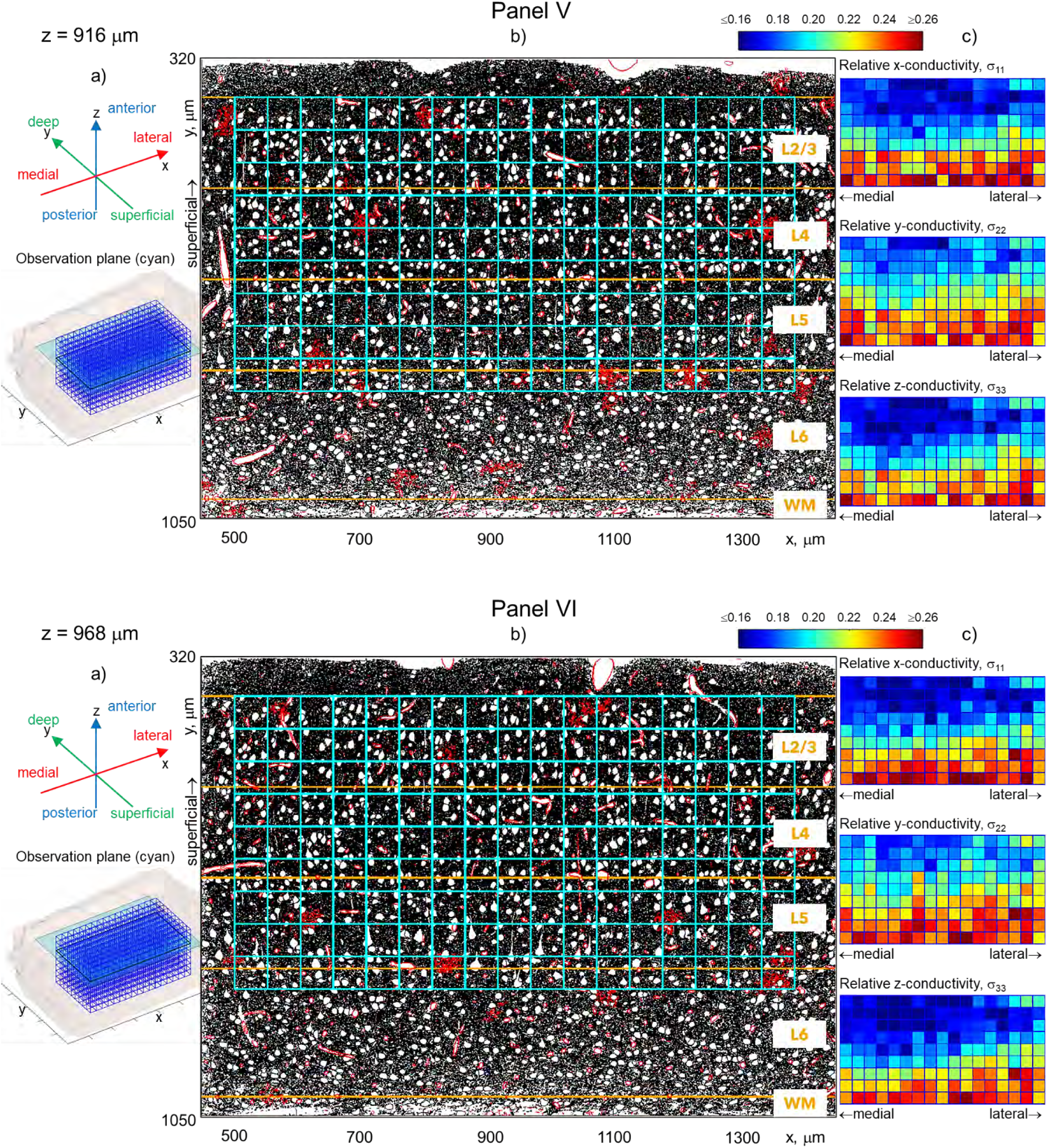
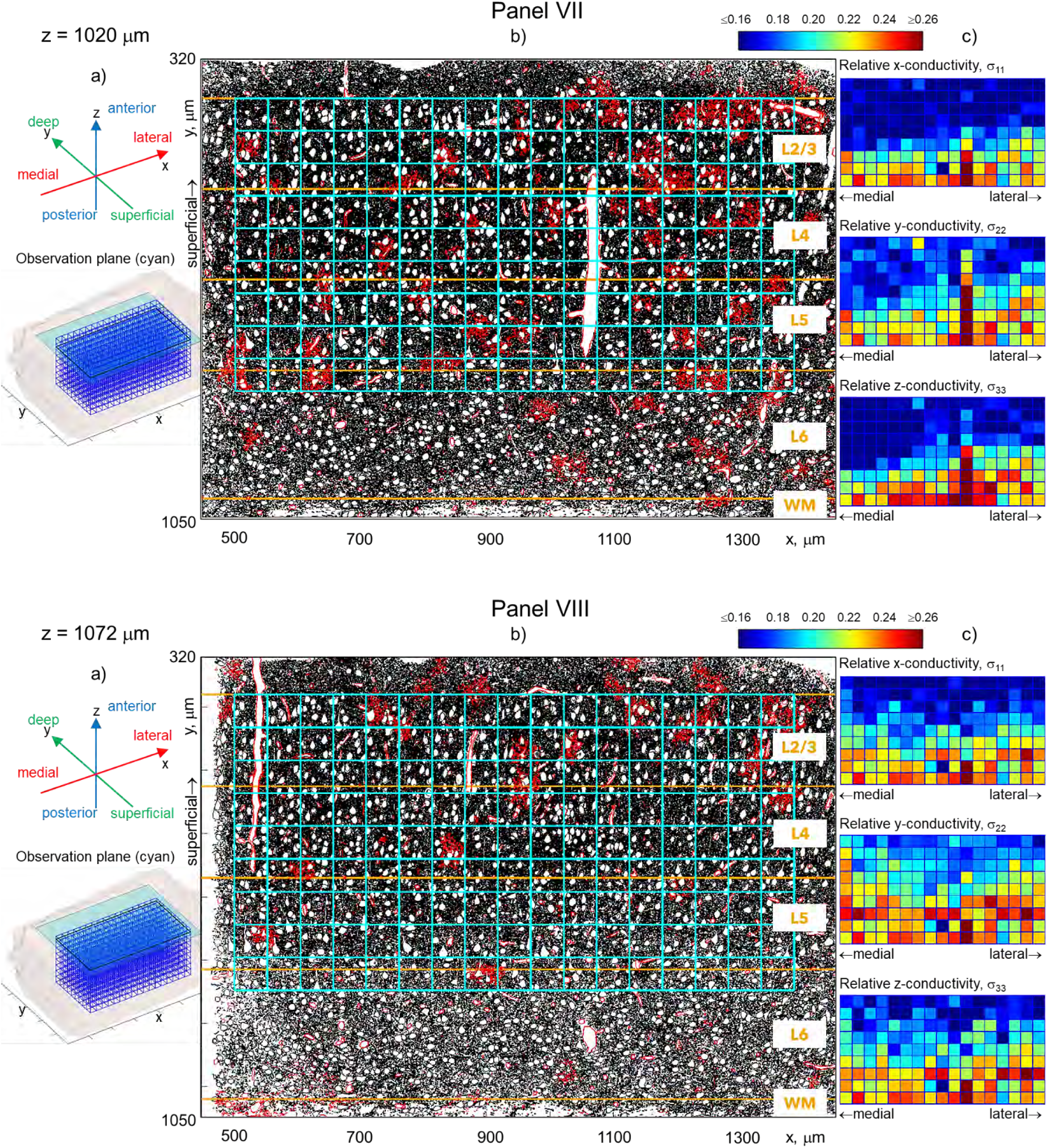
Panels I-VIII. Two-dimensional conductivity maps which gradually evolve from anterior to posterior direction over the total distance of approximately 360 μm. Panels I-VIII a) Position of the observation plane within the sample. Panels I-VIII b) Cross-section of the original *Minnie 65* cellular volume at the given value of the original *zz* -coordinate of the sample cross-section and containing the selected block grid in the coronal (frontal) plane. Here, objects in black correspond to cell membranes; objects in red indicate the blood network and astrocytes; white spots are somas of individual neurons. Block projections (blue) and boundaries between cortical layers (orange) are also shown there. Approximate layer boundaries (derived from 3D boundary meshes) shown are based on local nucleus features and are plotted following [40].; Panels I-VIII c) Three two-dimensional conductivity maps for three different components of the conductivity tensor for blocks whose centers coincide with the given cross-sectional coronal plane. Every such map comprises 17×9 blocks. Different conductivity values are encoded by different colors; they are again normalized to the conductivity of the homogeneous space filled with only extracellular fluid whose reference conductivity is set as 1 *SS*/*mm*. The same color scale, from 0.16 to 0.26, is used in all these panels, which is represented by a colorbar on top.

Fig. 6a (for every Panel I through VIII) displays movement of the cross-section plane from posterior to anterior within the sample. Fig. 6b (again for every Panel I-VIII) shows the corresponding cross-section of the original *Minnie 65* volume at the given value of the original *z* -coordinate of the sample. There, objects in black correspond to cell membranes; objects in red indicate the blood network of capillaries and astrocytes; white spots are somas of individual neurons. Block projections (blue) and boundaries between cortical layers (orange) are also shown. The approximate layer boundaries shown (derived from 3D boundary meshes) are based on local nucleus features and are plotted following [40].

Fig. 6c (for every Panel I through VIII) gives the three two-dimensional conductivity maps for blocks whose centers coincide with the given cross-sectional coronal plane. Each map comprises 17×9 blocks. Different conductivity values are encoded by different colors; they are again normalized to the conductivity of the homogeneous space filled with only extracellular fluid whose reference conductivity is set as 1 *SS*/*mm*. The same color scale, from 0.16 to 0.26, is used in all these figures, which is represented by a colorbar on top.

### 3.2 Three-dimensional conductivity maps

Fig. 7 displays anisotropic normalized voxel conductivity data for all 1,224 square blocks spanning L2/L3, L4, and L5 of the *Minnie 65* MICrONS sample shown by different colors of labeling spheres located at the center of each block. Fig. 7a is the conductivity in the medial-lateral direction, *σ*_11_; Fig. 7b is the conductivity in the deep-superficial direction, *σ*_22_; and Fig. 7c is the conductivity in the anterior-posterior direction, *σ*_33_. There, different cross-section planes correspond to different layer boundaries. Maximum and minimum conductivity values are also given for every principal direction.

**Fig. 7.**
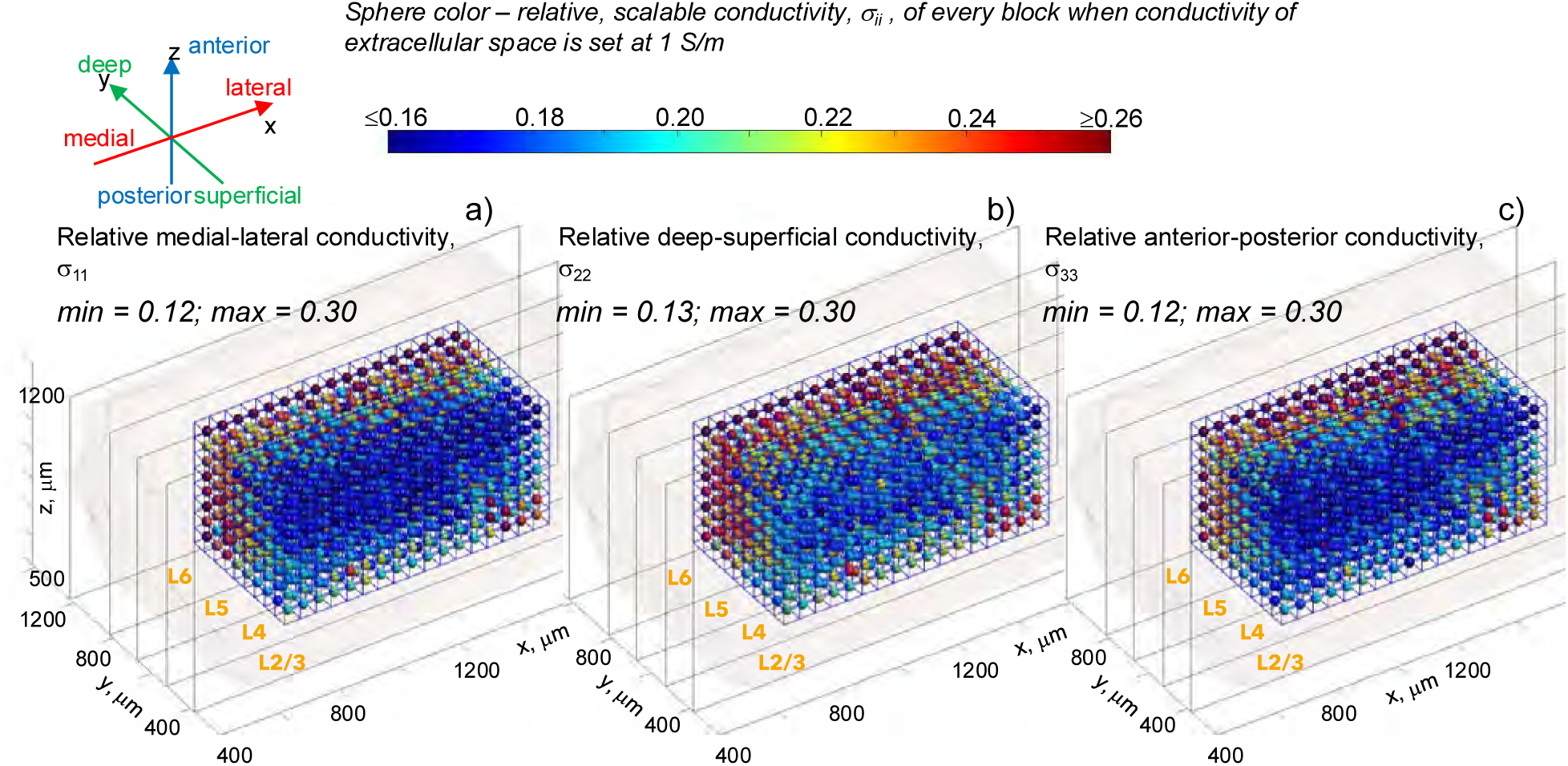
a-c) Anisotropic normalized voxel conductivity data for 1,224 square blocks spanning L2/L3, L4, and L5 of the *Minnie 65* MICrONS sample shown by different colors of labeling spheres located at the center of each block. a) Conductivity in the medial-lateral direction, *σ*_11_. b) Conductivity in the deep-superficial direction, *σ*_22_. c) Conductivity in the anterior-posterior direction, *σ*_33_. In each figure, different cross-section planes correspond to different layer boundaries. Maximum and minimum conductivity values are also given for every principal direction.

All conductivity values in Fig. 7 are normalized to that of the homogeneous extracellular space filled with only extracellular fluid whose reference conductivity is chosen as 1 *S*/*m*. To obtain absolute conductivity values, multiplication by the realistic conductivity of the extracellular fluid (the ionic solution filling the extracellular space or ECS), *σ*_ECS_, should be made. This extracellular conductivity – which is often associated with the conductivity of the extracellular fluid – is discussed in a number of sources [20],[21],[41],[42]. If, for example, *σ*_ECS_ = 1.5 *SS*/*mm* and the relative medial-lateral conductivity, *σ*_11_ of a block in Fig. 7a is 0.15 then the absolute medial-lateral conductivity value of the block should be obtained as 0.225 *S*/*m*. The numerical three-dimensional datasets for *σ*_11_, *σ*_22_, and *σ*_33_ are available as supplementary materials and online [43].

### 3.3 Average conductivity in three principal planes – evolution along and across the cortex

Fig. 8 reports average conductivity values when moving along and across the cortex, respectively, in the three principal planes and for the entire sample. Fig. 8a,b yields the normalized conductivity values averaged in the sagittal plane of the *Minnie 65* sample – when moving along the cortex. The plane is moving from medial to lateral direction as shown in Fig. 8a. The red curve is the conductivity in the medial-lateral direction, *σ*_11_; the green curve is the conductivity in the deep-superficial direction, *σ*_22_; and the blue curve is the conductivity in the anterior-posterior direction, *σ*_33_.

**Fig. 8.**
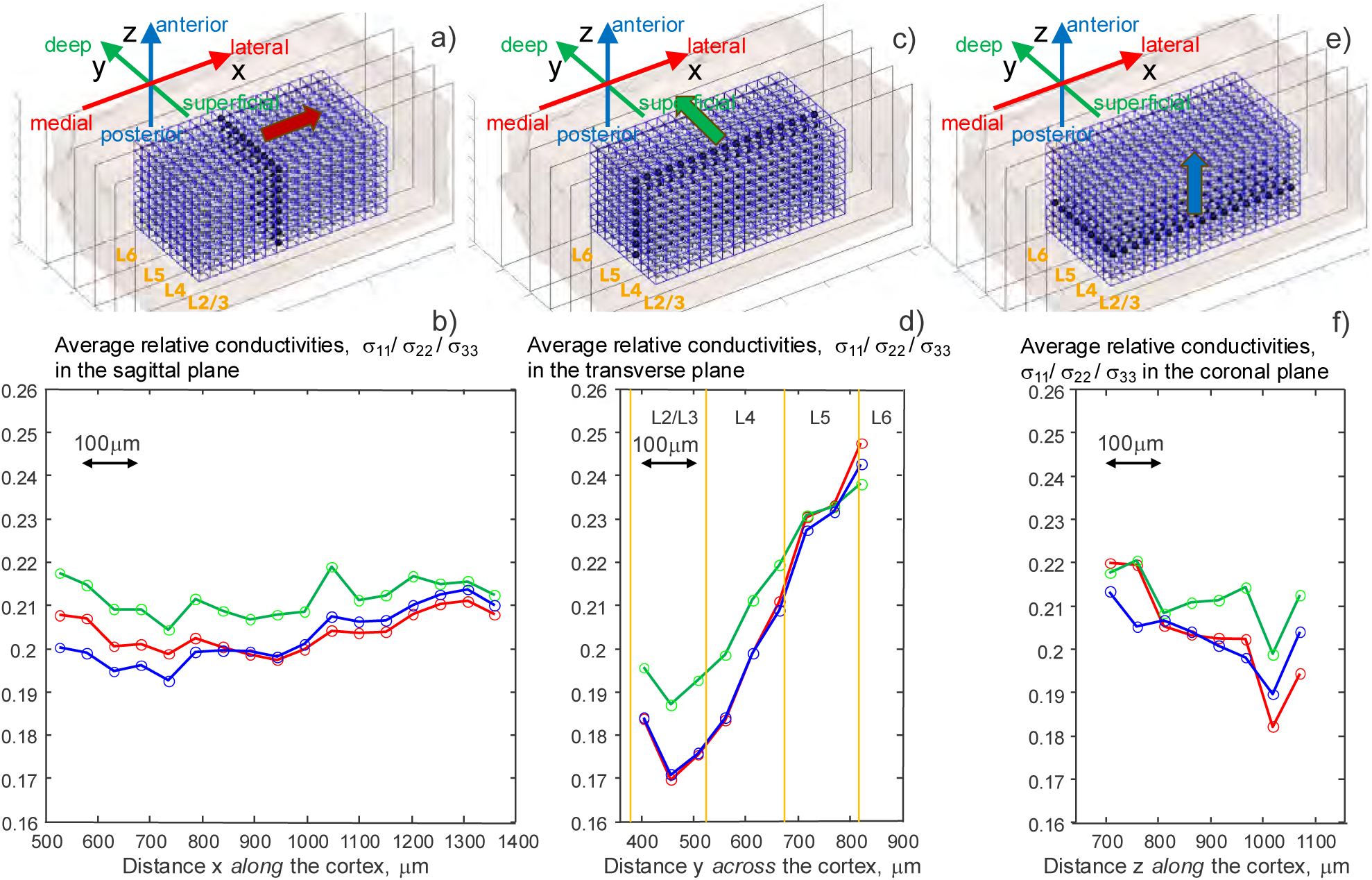
Average conductivity values when moving along and across the cortex, respectively, in the three principal planes. a,b) Normalized conductivity values averaged in the sagittal plane of the *Minnie 65* sample – when moving along the cortex. The plane is moving from medial to lateral direction as shown in a). Red curve is the conductivity in the medial-lateral direction, *σ*_11_; green curve is the conductivity in the deep-superficial direction, *σ*_22_; and blue curve is the conductivity in the anterior-posterior direction, *σ*_33_. c,d) Normalized conductivity values averaged in the transverse plane of the *Minnie 65* sample – when moving across the cortex. The plane is moving from superficial to deep direction as shown in c). Red curve is again the conductivity in the medial-lateral direction, *σ*_11_; green curve is the conductivity in the deep-superficial direction, *σ*_22_; and blue curve is the conductivity in the anterior-posterior direction, *σ*_33_. e,f) Normalized conductivity values averaged in the coronal plane of the *Minnie 65* sample – when again moving along the cortex. The plane is moving from posterior to anterior direction as shown in e). As before, the red curve is the conductivity in the medial-lateral direction, *σ*_11_; the green curve is the conductivity in the deep-superficial direction, *σ*_22_; and the blue curve is the conductivity in the anterior-posterior direction, *σ*_33_.

Fig. 8c,d gives normalized layer-based conductivity values averaged in the transverse plane of the *Minnie 65* sample – when moving across the cortex. The plane is moving in the direction from superficial to deep, as shown in Fig. 8c. Again, the red curve is the conductivity in the medial-lateral direction, *σ*_11_; the green curve is the conductivity in the deep-superficial direction, *σ*_22_; the blue curve is the conductivity in the anterior-posterior direction, *σσ*_33_.

Finally, Fig. 8e,f gives the normalized conductivity values averaged in the coronal plane of the *Minnie 65* sample when again moving along the cortex. The plane is now moving in the direction from posterior to anterior, as shown in Fig. 3e. As before, the red curve is the conductivity in the medial-lateral direction, *σ*_11_; the green curve is the conductivity in the deep-superficial direction, *σ*_22_; and the blue curve is the conductivity in the anterior-posterior direction, *σ*_33_. In Fig. 8, all three conductivity maps have identical spatial and conductivity scales.

### 3.4 Average conductivity in three principal planes for separate parts of the sample

Fig. 9 is conceptually similar to Fig. 8; however, averaging is now made not for the entire sample, but for two selected parts – either layers L2/L3/L4 only (Fig. 9a,b and Fig. 9e,f) or the medial part of the sample only (Fig. 9c,d) which comprises approximately half of the sample. The goal of this additional investigation is to show that the evolution of averaged radial (across the cortex) or tangential (along the cortex) conductivities remains quite similar both for the entire sample (Fig. 8) and for a macroscopic portion of it (Fig. 9). At the same time, an additional, substantial macroscale shift in the conductivity values may occur. It is especially well observed in Fig. 9a,b and in Fig. 9e,f.

**Fig. 9.**
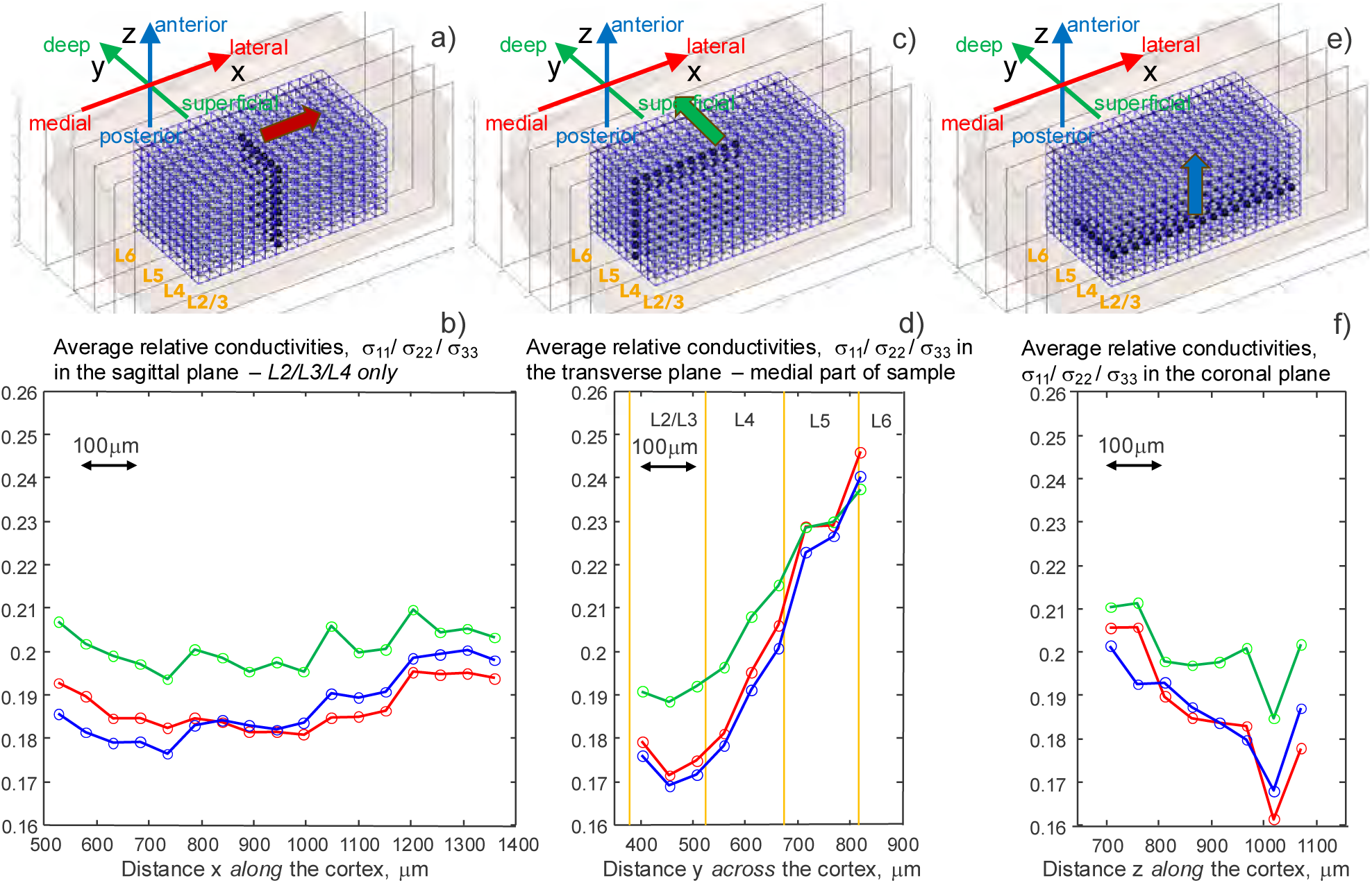
The same result as in Fig. 8, but averaging is now made not for the entire sample, but for two selected parts of it – either layers L2/L3/L4 only (a,b) or the medial part of the sample only (c,d) which comprises approximately a half of the sample. To be consistent with Fig 8, the three conductivity maps have the identical spatial scale as well the identical conductivity scale.

### 3.5 Granularity of gray matter conductivity

We evaluate average and local spatial granularity of the conductivity distribution using both the native 50-µm resolution and after coarse-graining to 100-µm resolution. The 100-µm scale is more important for its direct impact on neuronal activation. As the first metric, we consider exceedance or ‘at least’ probability *PP* which characterizes large relative conductivity differences – conductivity contrasts – between immediately neighboring blocks which are sharing the same face. For each face-sharing neighboring block pair with conductivities 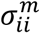 and 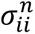, the symmetric relative difference defined as 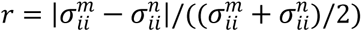. The at-least probability *P*(*r* ≥ *τ*) represents the probability that two neighboring blocks differ in conductivity by at least *τ*. Table 3 reports *P*(*r* ≥ *τ*) for *τ* = 5, 10, 20, 40, 80%.

**Table 3.**
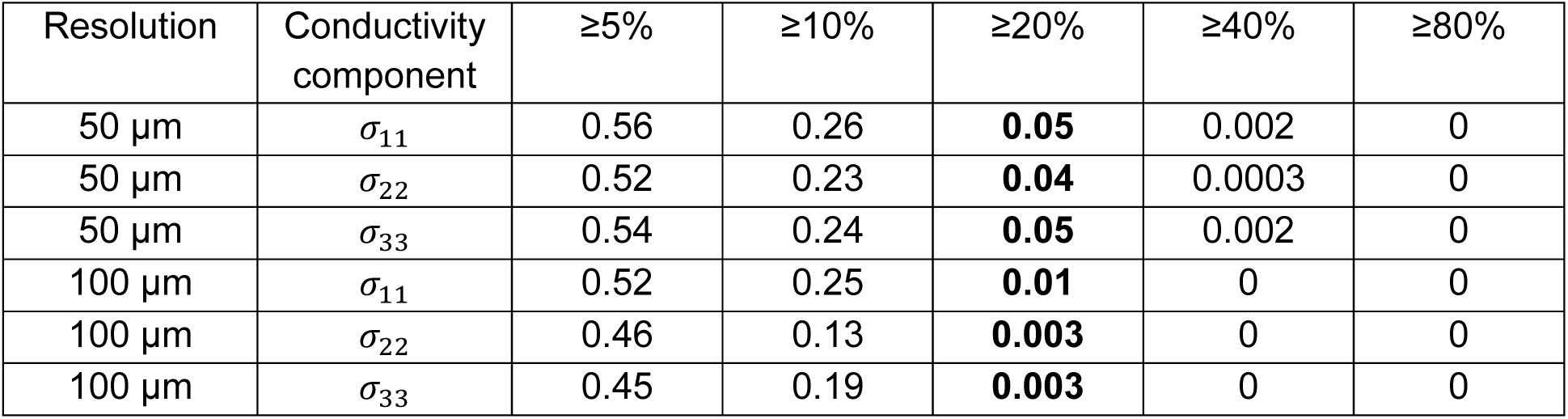
Probability *P* that two immediately neighboring gray matter blocks differ in conductivity by at least a relative amount *τ* for *τ* = 5, 10, 20, 40, 80%. Results are given at 50 µm resolution (3311 neighbor pairs per direction) and at 100 µm resolution after 2×2×2 averaging (304 neighbor pairs per direction).

| Resolution | Conductivity component | $\geq 5\%$ | $\geq 10\%$ | $\geq 20\%$ | $\geq 40\%$ | $\geq 80\%$ |
| --- | --- | --- | --- | --- | --- | --- |
| 50 $\mu\text{m}$ | $\sigma_{11}$ | 0.56 | 0.26 | **0.05** | 0.002 | 0 |
| 50 $\mu\text{m}$ | $\sigma_{22}$ | 0.52 | 0.23 | **0.04** | 0.0003 | 0 |
| 50 $\mu\text{m}$ | $\sigma_{33}$ | 0.54 | 0.24 | **0.05** | 0.002 | 0 |
| 100 $\mu\text{m}$ | $\sigma_{11}$ | 0.52 | 0.25 | **0.01** | 0 | 0 |
| 100 $\mu\text{m}$ | $\sigma_{22}$ | 0.46 | 0.13 | **0.003** | 0 | 0 |
| 100 $\mu\text{m}$ | $\sigma_{33}$ | 0.45 | 0.19 | **0.003** | 0 | 0 |

Further, two complementary global metrics were used: the coefficient of variation (CV) or global spread, which captures overall dispersion of conductivity values across all blocks, and a granularity index (GI) or global RMS roughness, which quantifies average brick-to-brick conductivity differences. The coefficient of variation, CV, is defined as the standard deviation of conductivity across blocks divided by the mean conductivity. The granularity index, GI, is defined as the root-mean-square conductivity difference between immediately neighboring blocks (six in total), normalized by mean conductivity. The results for CV and GI are collected in Table 4.

**Table 4.** Mesoscale heterogeneity and granularity (percentages) of normalized gray-matter conductivity at 50 µm and 100 µm resolution.

| Resolution | Conductivity component | Average CV, % | Average GI, % |
| --- | --- | --- | --- |
| 50 $\mu\text{m}$ | $\sigma_{11}$ | 16 | 9.3 |
| 50 $\mu\text{m}$ | $\sigma_{22}$ | 12 | 8.8 |
| 50 $\mu\text{m}$ | $\sigma_{33}$ | 15 | 8.9 |
| 100 $\mu\text{m}$ | $\sigma_{11}$ | 13 | 8.4 |
| 100 $\mu\text{m}$ | $\sigma_{22}$ | 9 | 6.7 |
| 100 $\mu\text{m}$ | $\sigma_{33}$ | 12 | 7.7 |

## 4. Discussion

### 4.1 Qualitative agreement with prior cortical conductivity data

The present study shows that

i. Spatially averaged electrical conductivity of the rodent’s cortical gray matter is lowest in L2/L3 and then gradually increases when progressing to L4 and further to L5 (Fig. 8d, Fig. 9d). This observation is in accordance with prior measurements [5],[8] as well as with indirect estimates [6] based on the amount of cortical extracellular space [20],[21]. Our conductivity variations across layers L2 through L5 for all three components of the conductivity tensor reach approximately 30% (for the radial conductivity *σσ*_22_) and 40% (for tangential conductivities *σ*_11_, *σ*_33_).
ii. Tangential conductivity (red and blue curves in Figs. 8,9) is always smaller than radial conductivity (green curves in Figs. 8,9) when progressing from L2/L3 to L5. This observation is also in accordance with prior measurements [5] (rats) and [44] (cats).

### 4.2 Quantitative agreement with prior cortical conductivity data

When compared to the biologically closest rodent’s case, the spatially averaged radial conductivity values, *σ*_22_, obtained in this study are in a good quantitative agreement with prior lower-resolution (300 µm) extensive multi-site LFP (local field potential) *in-vivo* recordings in rat somatosensory cortexes acquired using silicone microelectrode arrays [5]. To show this, Table 5 collects values from Fig. 8 averaged layerwise versus those of Ref. [5]. This presents not normalized but absolute average conductivity values and their margins layerwise for the commonly accepted ECS conductivity value of 1.79 S/m (cf., e.g., [1]).

**Table 5.** Absolute average conductivity values of the present study and their margins layerwise given the standard ECS conductivity of 1.79 S/m as compared to Ref [5].

| Layer | L2/L3 | L4 | L5 | L6 |
| --- | --- | --- | --- | --- |
| Conductivity across cortex (radial conductivity), present study, S/m | **0.34** | **0.38** | **0.42** | **NA** |
| Minimum block conductivity, S/m | 0.21 | 0.23 | 0.27 | **NA** |
| Maximum block conductivity, S/m | 0.42 | 0.50 | 0.53 | **NA** |
| Microelectrode-derived conductivity across cortex (radial conductivity), S/m – rat's somatosensory cortex (mean for 5 rats) [5] | **0.32** | **0.33** | **0.35** | **0.29** |

### 4.3 New observations

The present study indicates that

i. Cortical conductivity is granular at both 50-µm and 100-µm resolution (Fig. 6 and Tables 3,4). Approximately 4%–5% of neighboring block pairs exhibit relative conductivity differences of at least 20% at 50-µm resolution, while such large differences are rare but non-zero at 100-µm resolution (∼0.3–1.0% depending on orientation). A small number of blocks (23) at 50-µm resolution have the conductivity contrast of 50% or slightly higher relative to their nearest neighbors.
ii. Cortical conductivity changes not only in radial but also in the tangential direction. For the present sample, this effect is most pronounced in the coronal plane and for layers L2/L3 and L4 where the tangential conductivity variation may reach 30% (Fig. 9f).

These two factors may, at least in part, account for the large variations in cortical conductivity observed in Table 1. We might expect that even higher variations may be observed when larger brain samples (e.g., inclusion of *Minnie 35* along with *Minnie 65* – cf. Fig. 1a) will be studied.

### 4.4 Limitations of present methodology

The major limitation of the present study is in explicitly using secondary data to correct for the amount of extracellular space (Eqs. (2), (3)). This is because not all the cells have been included in the present analysis – the estimated unlabeled volume is between 10-20%. Despite this relatively small number, its effect on cortical conductivity is large since small variations in the insulating content of a densely packed cytoarchitecture produce first-order conductivity variations. We must emphasize that the *Minnie 65* repository [11],[12] does include an extra, smaller collection of segmented unlabeled cells, which contains new, missing glia cells, and other cell pieces. We have downloaded 18,089 such new meshes. However, we were unable to separate these properly and include new cells into the pipeline since they heavily overlap or intersect with multiple cells or cell pieces from the main repository. We believe that rapid advances in AI-assisted cellular segmentation will help us to overcome this technical difficulty and provide near-complete labeling of neuronal and non-neuronal cellular elements of the cortex.

A further limitation is the assumption of a non-conducting membrane, which restricts the present analysis to the quasi-static (near-DC) regime. This limitation can be addressed by formulating coupled interior and exterior Neumann problems linked through a complex membrane impedance. Such an extension would preserve the underlying BEM–FMM framework, but it will increase the computational cost by approximately a factor of four. This extension is reserved for future work.

In the present study, we modeled microcapillaries as open structures filled with the same extracellular fluid (Supplement, Fig. 2). This assumption is not strictly accurate. Although the conductivities of venous and arterial blood [2] are relatively high, they remain lower than that of the extracellular space. A correction of this simplification was evaluated for several blocks. It results in somewhat reduced conductivity variations in the vicinity of blood vessels (e.g., Fig. 6, Panel I) but does not alter the values reported in Tables 3 and 4 by more than 5–10%.

Finally, the living brain contains active elements that generate transmembrane currents associated with membrane potentials, ion channels, and synaptic activity. In general, the local current density in neural tissue might be expressed as ***J*** = *σ**E*** + ****J***_neuron_*, where ****J***_neuron_* represents neuronal current sources. Fortunately, ****J***_neuron_* is conservative; it starts and ends in the same neuron, which should be electrically neutral as a whole (the sum of inflowing/outflowing currents is zero), according, for example, to the familiar cable equation [45],[46],[47]. Therefore, neglecting localized, neuronal current sources of active neurons could be appropriate for the present, external electrode-based mesoscale analysis.

### 4.5 Advantages of present methodology

The conventional finite element method (FEM) exclusively used in brain stimulation models and other relevant computational neuroscience studies today [48],[49],[50],[51],[52][53],[54],[55] cannot model these dense, highly tortuous and large brain microstructures per se, as it fails at the initial step of generating a background tetrahedral mesh. Instead, either a macroscale model (an entirely structurally homogeneous cortical gray matter) [56],[57] or small-scale artificial cellular structures composed of spheres, cylinders, hexagons, etc. are presently used – cf. [54],[55],[58].

In contrast, the boundary element fast multipole method (BEM-FMM) overcomes this limitation [15],[16],[17],[18]. Built on the fast multipole algorithm – recognized as one of the greatest algorithmic discoveries of the 20th century [59],[60] – BEM-FMM can model tens of thousands of cell membranes and microcapillary walls. On a standard university server (512 GB of RAM, 64 cores), it handles membrane meshes with up to 100 million facets [18] and much larger problems when using partitioning of cortical volumes, as introduced in this study.

## 5. Conclusions

Electrical conductivity of cortical gray matter governs the magnitude and spatial distribution of electric fields generated by brain stimulation and intrinsic neuronal activity. However, reported macroscopic conductivity values vary significantly, limiting the fidelity of bioelectromagnetic models. In this study, we attempted to derive mesoscale conductivity maps of mouse visual cortex at 50-µm resolution directly from large-volume, segmented nanometer-scale electron microscopy data. Spatially averaged conductivity values predicted by our framework agree well with prior low-resolution measurements in rats. At the same time, we observed substantial conductivity granularity at 50-100 µm scales as well as significant conductivity variations in both radial and tangential directions. These results indicate that mesoscale conductivity heterogeneity could be an intrinsic structural property of the cortex. Limitations and extensions of this study have been discussed.

The major set of data generated in this study – the cortical conductivity maps for three components of the conductivity tensor of the MICrONS *Minnie 65* brain sample – are publicly available via a GitHub repository [43].

## 6. Acknowledgements

This study has received support from the NIH grants R01MH130490 and R01EB035484 (SNM), U24NS120053 (BD) as well as the Intramural Research Program of NIH, NIMH ZIAMH002955 (Z-DD), and Intramural Research Program of NIH, NIDA ZIADA000638 (HL). MB is supported by grants from Harold Shames and NIH: UG3DA048502, T34 GM137858, R01NS112996, R01NS101362, and T32GM136499. BR is supported by P41EB030006.

## 7. Author Contributions and Declaration of Interest

G.M.N. and S.N.M. conceived the study and designed the overall research framework. G.M.N., S.N.M., and G.C.N.P. developed and implemented the computational modeling approach, including large-scale boundary-element fast multipole simulations. G.M.N., A.M., D.T., M.I., C.P., R.L., and E.H.B. contributed to data processing, numerical analysis, and validation. B.D. provided expertise on electron microscopy datasets and contributed to interpretation of cortical microstructure. K.W. and T.R.K. contributed to methodological discussions and interpretation of conductivity anisotropy and spatial variability. Z.-D.D., H.L., B.R.R., and M.B. contributed to interpretation of the results in the context of brain stimulation, neuroimaging, and neurophysiology. G.M.N. M.B., and S.N.M. led data interpretation and wrote the manuscript. All authors contributed to discussion of the results and provided critical feedback on the manuscript.

The authors declare no competing interests.

## References

[1]. Saturnino GB, Puonti O, Nielsen JD, Antonenko D, Madsen KH, Thielscher A. SimNIBS 2.1: A Comprehensive Pipeline for Individualized Electric Field Modelling for Transcranial Brain Stimulation. 2019 Aug 28. In: Makarov S, Horner M, Noetscher G, editors. Brain and Human Body Modeling: Computational Human Modeling at EMBC 2018 [Internet]. Cham (CH): Springer; 2019. Chapter 1. PMID: 31725247.

[2]. IT’IS Foundation. Low-Frequency (Conductivity) Tissue Properties Database. IT’IS Virtual Population. 2025. Tissue-specific conductivity values for human gray matter and other tissues up to 1 MHz.

[3]. McCann H, Pisano G, Beltrachini L. “Variation in Reported Human Head Tissue Electrical Conductivity.” Brain Topography. 2019;32(5):825–858. doi:10.1007/s10548-019-00710-2.

[4]. Huang Y, Liu AA, Lafon B, Friedman D, Dayan M, Wang X, Bikson M, Doyle WK, Devinsky O, Parra LC. Measurements and models of electric fields in the in vivo human brain during transcranial electric stimulation. Elife. 2017 Feb 7;6:e18834. doi:10.7554/eLife.18834. PMID: 28169833; PMCID: PMC5370189.

[5]. Goto T, Hatanaka R, Ogawa T, Sumiyoshi A, Riera J, Kawashima R. An Evaluation of the Conductivity Profile in the Somatosensory Barrel Cortex of Wistar Rats. J Neurophysiol 2010;104:3388–412. 10.1152/jn.00122.2010.

[6]. Wang B, Worbs T, Hussain MA, Aberra AS, Thielscher A, Grill WM, Peterchev AV. Mesoscale tissue properties and electric fields in brain stimulation -bridging the macroscopic and microscopic scales. arXiv:2511. Oct. 2025, 16465. 10.48550/arXiv.2511.16465.

[7]. Jahng GH, Lee MB, Kim HJ, Je Woo E, Kwon OI. Low-frequency dominant electrical conductivity imaging of in vivo human brain using high-frequency conductivity at Larmor-frequency and spherical mean diffusivity without external injection current. Neuroimage. 2021 Jan 15;225:117466. doi: 10.1016/j.neuroimage.2020.117466. Epub 2020 Oct 16. PMID: 33075557.

[8]. Lee S, Falchier A, Opitz A. Layer-specific electric fields and effective conductivity in nonhuman primates during transcranial electrical stimulation. Brain Stimul. 2026 Feb 18;19(2):103060. doi: 10.1016/j.brs.2026.103060.

[9]. Lim MM, Gerstner JR, Holtzman DM. The sleep–wake cycle and Alzheimer’s disease: what do we know? Neurodegener Dis Manag. 2014;4(5):351–362. doi:10.2217/nmt.14.33. PMID: 25405649.

[10]. Xie L, Kang H, Xu Q, Chen MJ, Liao Y, Thiyagarajan M, O’Donnell J, Christensen DJ, Nicholson C, Iliff JJ, Takano T, Deane R, Nedergaard M. Sleep drives metabolite clearance from the adult brain. Science. 2013 Oct 18;342(6156):373–7. doi: 10.1126/science.1241224. PMID: 24136970; PMCID: PMC3880190.

[11]. MICrONS Consortium. Functional connectomics spanning multiple areas of mouse visual cortex. Nature. 2025; (special issue). doi:10.1038/s41586-025-08790-w.

[12]. MICrONS Explorer. Public Data Release V. 1621. Online: https://www.microns-explorer.org/

[13]. Turner NL, Macrina T, Bae JA, et al. Multiscale and multimodal reconstruction of cortical structure and function. Cell. 2022;185(6):1084–1103.e24. doi:10.1016/j.cell.2022.02.039.

[14]. Barlow JW, Turner NL, Macrina T, et al. A connectome and analysis of a mammalian cortical inhibitory neuron class. Neuron. 2023;111(5):747–762.e7. doi:10.1016/j.neuron.2022.12.026.

[15]. Makarov SN, Noetscher GM, Raij T, Nummenmaa A. A Quasi-Static Boundary Element Approach with Fast Multipole Acceleration for High-Resolution Bioelectromagnetic Models. IEEE Trans Biomed Eng. 2018 Mar 7. doi: 10.1109/TBME.2018.2813261.

[16]. Makaroff SN, Nummenmaa AR, Noetscher GM, Qi Z, McIntyre CC, Bingham CS. Influence of charges deposited on membranes of human hyperdirect pathway axons on depolarization during subthalamic deep brain stimulation. J Neural Eng. 2023 Jul 19;20(4). doi: 10.1088/1741-2552/ace5de.

[17]. Noetscher GM, Tang D, Nummenmaa AR, Bingham CS, McIntyre CC, Makaroff SN. Estimations of Charge Deposition onto Convoluted Axon Surfaces within Extracellular Electric Fields. IEEE Trans Biomed Eng. 2023 Aug 3; doi: 10.1109/TBME.2023.3299734 (selected as the best paper of the issue).

[18]. Qi Z, Noetscher GM, Miles A, Weise K, Knösche TR, Cadman CR, Potashinsky AR, Liu K, Wartman WA, Nunez Ponasso G, Bikson M, Lu H, Deng ZD, Nummenmaa AR, Makaroff SN. Enabling Electric Field Model of Microscopically Realistic Brain. Brain Stimul. 2025 Jan;18(1):77–93. doi: 10.1016/j.brs.2024.12.1192.

[19]. Shapson-Coe A, Januszewski M, Berger DR, Pope A, Wu Y, Blakely T, Schalek RL, Li PH, Wang S, Maitin-Shepard J, Karlupia N, Dorkenwald S, Sjostedt E, Leavitt L, Lee D, Troidl J, Collman F, Bailey L, Fitzmaurice A, Kar R, Field B, Wu H, Wagner-Carena J, Aley D, Lau J, Lin Z, Wei D, Pfister H, Peleg A, Jain V, Lichtman JW. A petavoxel fragment of human cerebral cortex reconstructed at nanoscale resolution. Science. 2024 May 10;384(6696):eadk4858. doi: 10.1126/science.adk4858.

[20]. Lehmenkühler A, Syková E, Svoboda J, Zilles K, Nicholson C. Extracellular space parameters in the rat neocortex and subcortical white matter during postnatal development determined by diffusion analysis. Neuroscience. 1993 Jul;55(2):339–51. doi: 10.1016/0306-4522(93)90503-8. PMID: 8377929.

[21]. Syková E, Nicholson C. Diffusion in brain extracellular space. Physiol Rev. 2008 Oct;88(4):1277–340. doi: 10.1152/physrev.00027.2007. PMID: 18923183; PMCID: PMC2785730.

[22]. Wan J, Li W, Adhinarta JK, Banerjee A, Sjostedt E, Wu J, Lichtman J, Pfister H, Wei D. TriSAM: Tri-Plane SAM for Zero-Shot Cortical Blood Vessel Segmentation in VEM Images. IEEE J Biomed Health Inform. 2025 Nov;29(11):8246–8255. doi: 10.1109/JBHI.2025.3577625. PMID: 40489282.

[23]. Alliez P, Cohen-Steiner D, Hemmer M, Portaneri C, Rouxel-Labbé M. 3D Alpha Wrapping. In CGAL User and Reference Manual. CGAL Editorial Board, 5.6 edition, 2023.

[24]. Korogod N, Petersen CC, Knott GW. Ultrastructural analysis of adult mouse neocortex comparing aldehyde perfusion with cryo fixation. Elife. 2015 Aug 11;4:e05793. doi: 10.7554/eLife.05793. PMID: 26259873; PMCID: PMC4530226.

[25]. Hama K, Arii T, Edamatsu M, et al. Measurement of the extracellular space in the rat brain cortex by diffusion-weighted magnetic resonance imaging. J Neurosci. 2004 Nov 3;24(44):9596–9600. doi:10.1523/JNEUROSCI.2639-04.2004. PMID: 15509750.

[26]. Hildebrand DGC, Cicconet M, Torres RM, Choi W, Quan TM, Moon J, Wetzel AW, Scott Champion A, Graham BJ, Randlett O, Plummer GS, Portugues R, Bianco IH, Saalfeld S, Baden AD, Lillaney K, Burns R, Vogelstein JT, Schier AF, Lee WA, Jeong WK, Lichtman JW, Engert F. Whole-brain serial-section electron microscopy in larval zebrafish. Nature. 2017 May 18;545(7654):345–349. doi: 10.1038/nature22356.

[27]. Hashin Z, Shtrikman S. A variational approach to the theory of the effective magnetic permeability of multiphase materials. J Appl Phys. 1962;33:3125–3131.

[28]. Landauer R. Electrical conductivity in inhomogeneous media. AIP Conf Proc. 1978;40:2–45.

[29]. Maxwell Garnett JC. Colours in metal glasses and metal films. Philos Trans R Soc Lond A. 1904;203:385–420.

[30]. Basser PJ, Roth BJ. The electrical conductivity of extracellular media in the brain. Biophys J. 1991;59:1147–1159.

[31]. Koch C. Biophysics of computation: information processing in single neurons. New York: Oxford University Press; 1999.

[32]. Destexhe A, Rudolph M, Paré D. The high-conductance state of neocortical neurons in vivo. Nat Rev Neurosci. 2003;4(9):739–751. doi:10.1038/nrn1198. PMID:12951566.

[33]. Markram H, Muller E, Ramaswamy S, Reimann MW, Abdellah M, Sanchez CA, et al. Reconstruction and simulation of neocortical microcircuitry. Cell. 2015;163(2):456–492. doi:10.1016/j.cell.2015.09.029. PMID:26451489.

[34]. Gentet LJ, Stuart GJ, Clements JD. Direct measurement of specific membrane capacitance in neurons. Biophys J. 2000;79(1):314–320. doi:10.1016/S0006-3495(00)76287-8. PMID:10866957.

[35]. Krassowska W, Neu JC. Response of a single cell to an external electric field. Biophys J. 1994 Jun;66(6):1768–76. doi: 10.1016/S0006-3495(94)80971-3.

[36]. Klee M, Plonsey R. Stimulation of spheroidal cells--the role of cell shape. IEEE Trans Biomed Eng. 1976 Jul;23(4):347–54. doi: 10.1109/tbme.1976.324597.

[37]. Cartee LA, Plonsey R. The transient subthreshold response of spherical and cylindrical cell models to extracellular stimulation. IEEE Trans Biomed Eng. 1992 Jan;39(1):76–85. doi: 10.1109/10.108130.

[38]. Greengard L, Gueyffier D, Martinsson P-G, Rokhlin V. Fast direct solvers for integral equations in complex three-dimensional domains. Acta Numer. 2009 May;18:243–275. doi:10.1017/S0962492906410011.

[39]. Flatiron Institute Fast Multipole Libraries. fmm3D Documentation Nov. 2024. Online: https://github.com/flatironinstitute/FMM3D.

[40]. Online: https://spelunker.cave-explorer.org/#!middleauth+https://global.daf-apis.com/nglstate/api/v1/5901681641390080 (perspective view at z: 770). Accessed April 2025.

[41]. Neuroglancer -WebGL-based viewer for volumetric data capable of interactive visualization of cross-sections, meshes, and skeletons. Online: https://github.com/google/neuroglancer.

[42]. Baumann SB, Wozny DR, Kelly SK, Meno FM. The electrical conductivity of human cerebrospinal fluid at body temperature. IEEE Trans Biomed Eng. 1997 Mar;44(3):220–3. doi: 10.1109/10.554770. PMID: 9216137.

[43]. IT’IS Foundation Database at low frequencies. Online: https://itis.swiss/virtual-population/tissue-properties/database/low-frequency-conductivity/?utm_source=chatgpt.com

[44]. Tensor Conductivity Maps – ‘Mesoscale Cortical Conductivity at 50 µm Resolution’. 02/02/2026.GitHub repository: https://github.com/dexuantang/Mesoscale-Cortical-Conductivity-at-50-m-Resolution.

[45]. Hoeltzell PB, Dykes RW. Conductivity in the somatosensory cortex of the cat --evidence for cortical anisotropy. Brain Res. 1979 Nov 9;177(1):61–82. doi: 10.1016/0006-8993(79)90918-1.

[46]. Rattay F. Analysis of models for external stimulation of axons. IEEE Trans Biomed Eng. 1986 Oct;33(10):974–7. doi: 10.1109/TBME.1986.325670. PMID: 3770787.

[47]. Rattay F. Modeling the excitation of fibers under surface electrodes. IEEE Trans Biomed Eng. 1988 Mar;35(3):199–202. doi: 10.1109/10.1362.

[48]. Rattay F. Analysis of models for extracellular fiber stimulation. IEEE Trans Biomed Eng. 1989 Jul;36(7):676–82. doi: 10.1109/10.32099.

[49]. Seo H, Jun SC. Multi-Scale Computational Models for Electrical Brain Stimulation. Front Hum Neurosci. 2017 Oct 26;11:515. doi: 10.3389/fnhum.2017.00515. Review.

[50]. Aberra AS, Peterchev AV, Grill WM. Biophysically realistic neuron models for simulation of cortical stimulation. J Neural Eng. 2018 Dec;15(6):066023. doi: 10.1088/1741-2552/aadbb1.

[51]. Aberra AS, Wang B, Grill WM, Peterchev AV. Simulation of transcranial magnetic stimulation in head model with morphologically-realistic cortical neurons. Brain Stimul. 2020 Jan-Feb;13(1):175–189. doi: 10.1016/j.brs.2019.10.002.

[52]. Shirinpour S, Hananeia N, Rosado J, Tran H, Galanis C, Vlachos A, Jedlicka P, Queisser G, Opitz A. Multi-scale modeling toolbox for single neuron and subcellular activity under Transcranial Magnetic Stimulation. Brain Stimul. 2021 Sep 22:S1935-861X(21)00234–5. doi: 10.1016/j.brs.2021.09.004.

[53]. Menon P, Pavey N, Aberra AS, van den Bos MAJ, Wang R, Kiernan MC, Peterchev AV, Vucic S. Dependence of cortical neuronal strength-duration properties on TMS pulse shape. Clin Neurophysiol. 2023 Jun;150:106–118. doi: 10.1016/j.clinph.2023.03.012.

[54]. Konstantin Weise, Torge Worbs, Benjamin Kalloch, Victor H. Souza, Aurélien Tristan Jaquier, Werner Van Geit, Axel Thielscher, Thomas R. Knösche; Directional sensitivity of cortical neurons towards TMS-induced electric fields. Imaging Neuroscience 2023; 1 1–22. doi: 10.1162/imag_a_00036.

[55]. Wang B, Aberra AS. Bridging macroscopic and microscopic modeling of electric field by brain stimulation. Brain Stimul. 2025 May-Jun;18(3):897–899. doi: 10.1016/j.brs.2025.04.009.

[56]. Wang B, Worbs T, Hussain MA, Aberra AS, Thielscher A, Grill WM, Peterchev AV. Mesoscale tissue properties and electric fields in brain stimulation -bridging the macroscopic and microscopic scales. arXiv:2511. Oct. 2025, 16465. 10.48550/arXiv.2511.16465.

[57]. Aberra AS, Wang B, Grill WM, Peterchev AV. Simulation of transcranial magnetic stimulation in head model with morphologically-realistic cortical neurons. Brain Stimul. 2020 Jan-Feb;13(1):175–189. doi: 10.1016/j.brs.2019.10.002.

[58]. Weise K, Worbs T, Kalloch B, Souza VH, Jaquier AT, Van Geit W, Thielscher A, Knösche TR. Directional sensitivity of cortical neurons towards TMS-induced electric fields. Imaging Neuroscience 2023; 1 1–22. doi: 10.1162/imag_a_00036.

[59]. Ying W, Henriquez CS. Hybrid finite element method for describing the electrical response of biological cells to applied fields. IEEE Trans Biomed Eng. 2007;54(4):611–620. doi:10.1109/TBME.2006.889172.

[60]. Dongarra, J., & Sullivan, F. (2000). Guest Editors’ Introduction: The Top 10 Algorithms. Computing in Science & Engineering, 2(1), 22–23. doi: 10.1109/5992.841783.

[61]. Cipra, B. A. (2000). The Best of the 20th Century: Editors Name Top 10 Algorithms. SIAM News, 33(4), 1–2.

